# Identify Non-Mutational p53 Functional Deficiency in Human Cancers

**DOI:** 10.1101/2022.07.28.501874

**Authors:** Qianpeng Li, Yang Zhang, Sicheng Luo, Zhang Zhang, Ann L. Oberg, David E. Kozono, Hua Lu, Jann N. Sarkaria, Lina Ma, Liguo Wang

**Author notes:** Equal contribution (Qianpeng Li, Yang Zhang). Corresponding authors. (Wang L), (Ma L).

## Abstract

An accurate assessment of *TP53*’s functional status is critical for cancer genomic medicine. However, there is a significant challenge in identifying tumors with non-mutational p53 inactivations that are not detectable through DNA sequencing. These undetected cases are often misclassified as p53-normal, leading to inaccurate prognosis and downstream association analyses. To address this issue, we build the support vector machine (SVM) models to systematically reassess p53’s functional status in *TP53* wild-type (*TP53*^WT^) tumors from multiple TCGA cohorts. Cross-validation demonstrates the excellent performance of the SVM models with a mean AUC of 0.9822, precision of 0.9747, and recall of 0.9784. Our study reveals that a significant proportion (87-99%) of *TP53*^WT^ tumors actually have compromised p53 function. Additional analyses uncovered that these genetically intact but functionally impaired (termed as predictively reduced function of p53 or *TP53*^WT^-pRF) tumors exhibit genomic and pathophysiologic features akin to p53 mutant tumors: heightened genomic instability and elevated levels of hypoxia. Clinically, patients with *TP53*^WT^-pRF tumors experience significantly shortened overall survival or progression-free survival compared to those with *TP53*^WT^-pN (predictive normal function of p53) tumors, and these patients also display increased sensitivity to platinum-based chemotherapy and radiation therapy.

## Introduction

Genetic alterations resulting in the gain or loss of gene function are major drivers of cancer development, and the identification of genomic variants through tumor DNA sequencing has significantly advanced our understanding of cancer genetics [1-3]. The most representative example is tumor suppressor p53 (encoded by the *TP53* gene), which is a transcription factor that plays critical roles in preventing tumorigenesis and tumor progression [4-6]. *TP53* is the most frequently mutated gene that undergoes genetic inactivation in at least 20 different types of cancer [7]. Somatic mutation frequencies of *TP53* exceed 50% in ovarian, esophageal, pancreatic, lung, colorectal, uterine, head and neck, oral (gingivobuccal), soft tissue (leiomyosarcoma), gastric and biliary tract cancers (ICGC: https://dcc.icgc.org/genes/ENSG00000141510/mutations). As a result, *TP53* is one of the most extensively studied genes, and its transcriptional targets are well characterized [8]. The functional activities of p53 have been frequently associated with the response to chemo- and radiation therapy [9-14]. Moreover, p53 dysfunction has been linked to immunosuppression and immune evasion [15-17]. Therefore, accurately assessing the functional status of p53 is critical for prognosis and personalized medicine.

However, genetic alteration is not the sole mechanism responsible for disrupting or activating protein function; there is a growing recognition that p53 function can be inactivated even in tumors with genetically wild-type *TP53*. While sporadic studies have provided some support for this notion [18-24], a systematic evaluation using large clinical samples is lacking. On the other hand, the underlying mechanisms may involve various non-mutational factors such as the post-translational modification [25-29], DNA methylation [30], chromatin states [31], and interplaying with microRNAs [32, 33]. These molecular mechanisms are widespread, but their heterogeneous nature presents a formidable challenge in characterizing them. Hence, a reevaluation of p53 inactivation that goes beyond *TP53* mutations not only provides a more holistic comprehension of p53’s involvement in cancer biology but also paves the way for the development of p53-targeted therapies.

In this study, our objective is to re-assess the functional status of p53 in tumors with wild-type *TP53.* We hypothesized that both mutational and non-mutational inactivation of p53 could be reflected by the altered expression of genes regulated by p53. Toward this end, we first defined cancer type-specific gene sets (encompassing both direct and indirect targets of p53), which can serve as indicators of p53’s functional status. This was accomplished through a systematic literature review and differential expression analysis (DEA) of RNA-seq data. We then calculated the composite expression score (CES) derived from these genes using various algorithms, including GSVA (gene set variation analysis) [34], ssGSEA (single sample gene set enrichment analysis) [35], combined Z-score [36, 37], and the first principal component of PCA (principal component analysis). Next, we trained and validated SVM models using CESs from non-cancerous normal tissues (referred to as the “NT” group, assumed to have normal p53 function) and tumor samples harboring *TP53* truncating mutations (referred to as the “TM” group, assumed to have lost or reduced p53’s tumor suppressor function). These SVM models were applied to tumors with *TP53* wildtype (referred to as the “WT” group) and tumors with missense mutations (referred to as the “MM” group). Finally, we systematically evaluated our predictions using multi-omics and clinical data.

## Results

### Identification of cancer-specific gene sets reflecting p53’s functional status

We identified a set of p53-regulated genes whose expression levels are statistically associated with p53 truncation for each cancer type and used them as features to predict the functional status of p53 status in these cancers. First, we compiled a list of 147 genes, comprising both direct and indirect targets, from a comprehensive study conducted by Fischer [8]. We examined the expression profiles of these 147 genes using RNA-seq from the GTEx datasets, and assessed the p53 bindings using the ReMap2020 ChIP-seq database [38] (**Figure 1 a**). These genes were expressed across various normal tissues, and their promoters displayed evidence of p53 binding (Supplementary Table 1). To further validate these p53-regulated genes, we reanalyzed RNA-seq and ChIP-seq data obtained from MCF-7 cells that possess wild-type *TP53*, both before and after p53 activation by gamma irradiation. We observed minimal or weak p53 bindings at the promoters of p53-regulated genes (such as *ATF3*, *BTG2*) prior to p53 activation. However, upon p53 activation, significant p53 bindings were observed, accompanied by consistent upregulation in the expression levels of these genes [39] (Supplementary Figure 1).

**Figure 1.**
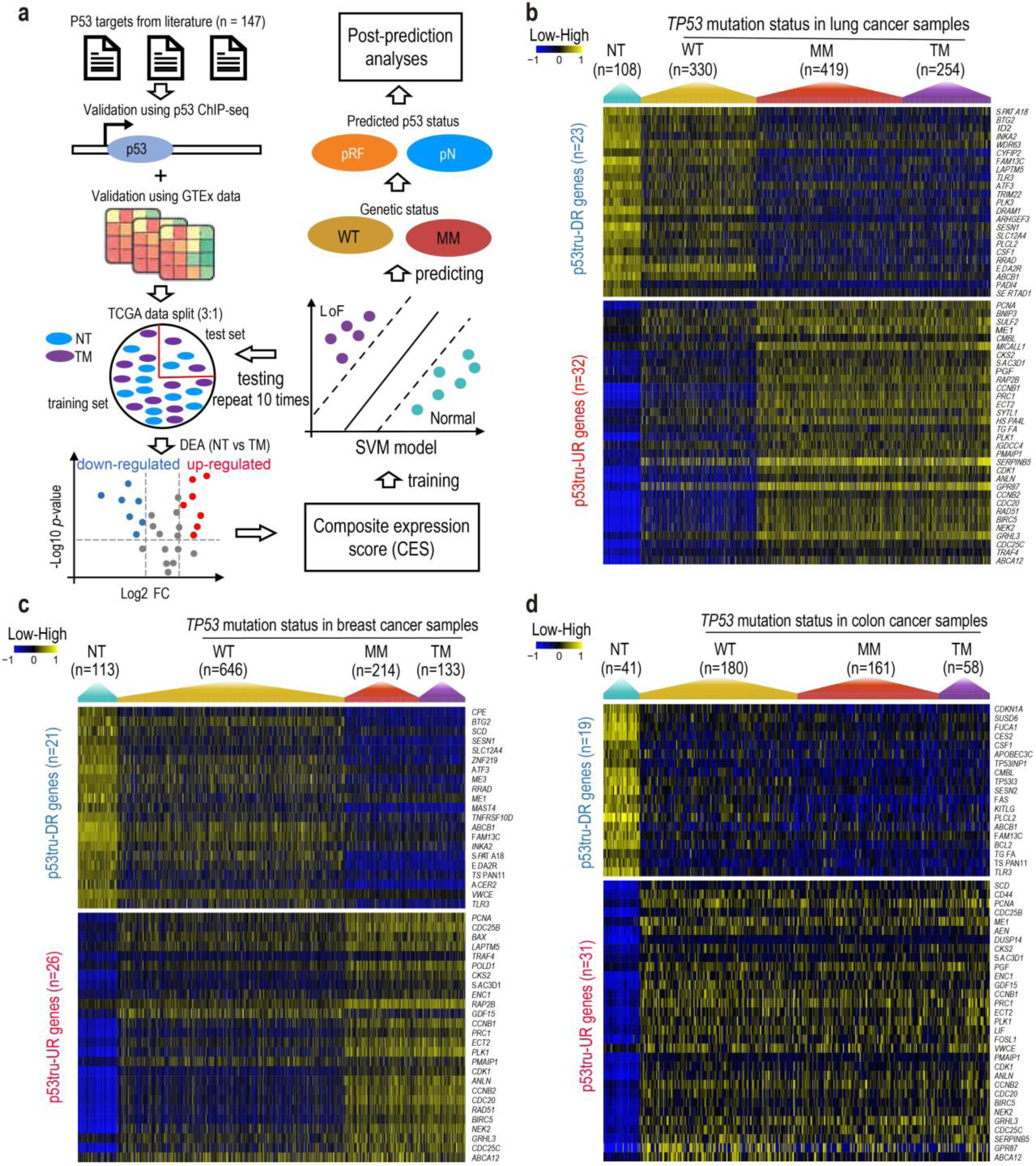
Identification of p53-regulated genes and their expression profiles in TCGA lung, breast, and colon cancers. (**a**) Analytic workflow depicting the process to identify p53tru-DR genes (genes down-regulated upon p53 truncation) and p53tru-UR genes (genes up-regulated upon p53 truncation), train and validate the SVM models. (**b**-**d**) Heatmaps illustrating the expression profiles of p53tru-DR and p53tru-UR genes in TCGA lung, breast, and colon cancers. The samples within each cohort were categorized into four groups based on TCGA designation: NT (adjacent normal tissue), WT (tumor tissue with wild-type *TP53*), MM (tumor tissue with *TP53* missense mutation or in-frame mutation), and TM (tumor tissue with *TP53* truncating mutation). CES, composite expression score; DEA, differential expression analysis; FC, fold change; GTEx, Genotype-Tissue Expression; pRF, predicted reduced function; pN, predicted normal; SVM, support vector machine.

The transcriptional programs regulated by p53 vary by tissue [40-42]. To define a gene set that can serve as features for predicting p53’s functional status in each selected cancer type, we conducted gene expression analyses between NT and TM groups. This analysis was performed using TCGA RNA-seq data for the previously selected 147 genes in each cancer type (**Figure 1 a**). As a result, we identified a set of genes that were down-regulated (referred to as p53tru-DR genes) and a set of genes that were up-regulated (referred to as p53tru-UR genes) in the TM group for each specific cancer type. It should be noted that p53tru-UR genes primarily represent genes whose expression is negatively regulated by normal p53, and the underlying mechanisms are likely indirect and not yet fully understood [8]. Finally, we identified 55, 47, 50, 39, 48, 58, and 50 genes regulated by p53 for TCGA lung cancer (LUNG), breast cancer (BRCA), colon cancer (COAD), head and neck cancer (HNSC), stomach cancer (STAD), endometrioid cancer (UCEC) and liver cancer (LIHC) cohorts, respectively. Among them, 12 genes were shared across all seven cancer types (Supplementary Table 1). It is worth noting that we focused on identifying p53-regulated genes and building SVM models for these seven TCGA cancer types, because each of them has sufficient non-cancer normal samples (n > 30) and *TP53* truncating mutation samples (n > 30) for training and testing. Additionally, we reanalyzed an independent lung adenocarcinoma (LUAD) dataset of East Asians to verify the selected gene set (see Materials and Methods). We identified 40 (out of 147 candidates) p53-regulated genes in this dataset using the same method. First, a high degree (37 or 92.5%) of the genes overlapped with p53-regulated genes identified in the TCGA LUNG cohort, and showed consistent up- and down-regulated direction. Second, we used p53-regulated genes identified in the TCGA LUNG cohort and performed unsupervised clustering to assess their potential representation of the p53 functional status (NT and TM) in this East Asian dataset. We found that all of the TM samples were predicted to be pRF, and 98.8% of the NT samples were predicted to be pN. These results demonstrate the robustness of our gene selection strategy.

### Characterizing composite expression of p53tru-DR and p53tru-UR genes

We analyzed the expression profiles of p53tru-DR and p53tru-UR genes across four distinct groups: NT, WT, MM, and TM in TCGA LUNG, BRCA, and COAD cohorts. As anticipated, p53tru-DR and p53tru-UR genes exhibited the opposite trends; the expressions of p53tru-DR genes were significantly decreased in the TM group, while it was significantly increased in the TM group, which is consistent with the compromised p53 status in this group (**Figure 1 b-d**). Interestingly, similar expression patterns of these p53-regulated genes were observed in the MM group (**Figure 1 b-d**). This suggests impaired p53 function in the MM samples, and indicates that missense/in-frame mutations and truncating mutations have a comparable impact on p53’s cellular activity. This observation aligns with the fact that most missense mutations occur within the DNA binding domain, leading to the disruption of p53’s ability to bind to DNA and transactivate its downstream targets [43]. These findings collectively indicated a reduced tumor suppressor function of p53 in *TP53*^WT^ samples.

Our hypothesis is that the impaired p53 function could be predicted by the altered expression of its regulated genes, regardless of its mutational status. To characterize the overall impact of a set of genes, we calculated the composite expression score (CES) of p53tru-DR and p53tru-UR genes using four different algorithms, including GSVA [34], ssGSEA [35], Z-score [36, 37], and PCA (Supplementary Table 2). Compared to the expression level of individual genes, the CES not only provides a combined and stable measure of p53 activity but also reduces the dimensionality of the expression data and helps mitigate potential overfitting of the SVM model.

Using the LUNG dataset as an example, we observed significant inverse correlations between CESs calculated from p53tru-DR and p53tru-UR genes. The Pearson’s correlation coefficients were - 0.996, -0.998, and -0.749 for GSVA, ssGSEA, and combined Z-score respectively (Supplementary Figure 2 a-c), indicating that both p53tru-DR and p53tru-UR genes could equally reflect p53’s functional status. In addition, we observed significant positive correlations (Pearson’s correlation coefficients ranged from 0.874 to 0.996) among CESs calculated by different algorithms, indicating excellent concordances among these algorithms (Supplementary Figure 2 d-i). When comparing the CESs across the NT, WT, MM, and TM groups, we found CESs of the NT samples tightly congregated within a narrow range. In contrast, CESs of the tumor samples (including WT, MM, and TM groups) showed a higher degree of dispersion, suggesting the increased heterogeneity in p53 activity among tumor samples (Supplementary Figure 3). Consistent with the gene-level expression data (**Figure 1 b**), CESs of WT samples were intermediate between those of NT and TM groups, while the CESs of the MM samples resembled those of the TM samples (**Figure 2 a-j**).

**Figure 2.**
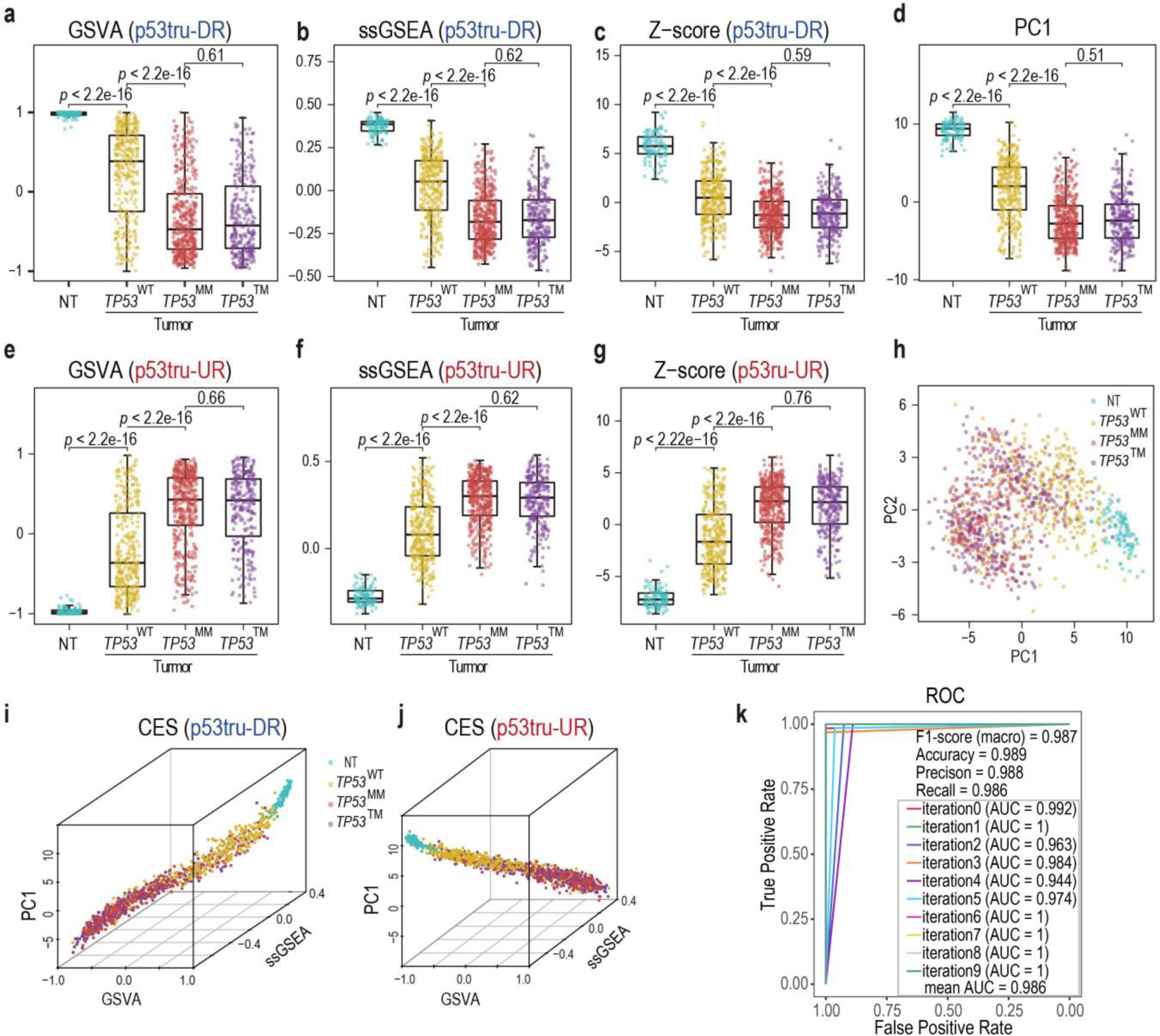
Composite expression scores (CESs) and the ROC curve of the SVM model built from the TCGA lung cancer cohort. (**a**-**c**) CESs of p53tru-DR genes calculated by GSVA, ssGSEA, and Z-score methods. In each panel, the blue, yellow, red, and purple box-and-whisker plots indicate the distribution of CESs in NT, WT, MM, and TM groups, respectively. (**d**) CES calculated by the first principal component scores (PC1) combining p53tru-DR and p53tru-UR genes. (**e-g**) CESs of p53tru-UR genes calculated by GSVA, ssGSEA, and Z-score methods. (**h**) Two-dimensional PCA plot demonstrating the clusters of NT, WT, MM, and TM samples. (**i-j**) Three-dimensional scatter plots illustrating the combinatorial effects of CES calculated from p53tru-DR genes and p53tru-UR genes, respectively. (**k**) Performance evaluation of the SVM model using ten-time hold-out validation. GSVA, gene set variation analysis; ssGSEA, single-sample gene set enrichment analysis; PCA, principal component analysis; AUC, area under the curve; ROC, receiver operating characteristic; NT, adjacent normal tissue; *TP53*^WT^, samples with wild-type *TP53*; *TP53*^MM^, samples with *TP53* missense mutation or in-frame mutation; *TP53*^TM^, samples with *TP53* truncating mutation; p53tru-DR genes, genes down-regulated upon p53 truncation; p53tru-UR genes, genes up-regulated upon p53 truncation.

### Building SVM models to predict p53 functional status in different cancer types

We proceeded to train SVM models using seven CESs calculated using four different algorithms, with the purpose of predicting p53’s functional status in *TP53*^WT^ and *TP53*^MM^ tumors. To achieve the best performance, these models were independently trained and validated for each cancer type (see Methods). Our cross-validation results exhibited outstanding performance by these SVM models. As exemplified by the LUNG cohort, the mean precision, recall, F1-score, and AUROC (area under the receiver operating characteristic curve) were 0.988, 0.986, 0.987, and 0.986, respectively (**Figure 2 k**). Similar performance was achieved in the remaining six cancer types (Supplementary Table 3).

The limited number of normal or truncating samples in other TCGA cancer types prevented us from training separate SVM models for these cancer types. Instead, we created a pan-cancer cohort and trained an SVM model by combining nine cancer types (LUNG, BRCA, COAD, HNSC, STAD, UCEC, LIHC, BLCA, ESCA; n = 5,160) with a relatively higher number of normal samples (n > 10) and truncating samples (n > 30) (Supplementary Figure 4). Despite the intrinsic heterogeneity of different cancer types, the pan-cancer SVM model demonstrated outstanding performance (Supplementary Figure 4 b, Supplementary Table 3). However, it should be noted that building a pan-caner SMV model is a compromise approach when individual cancer types lack sufficient samples to train their own SVM models, as it might be dominated by a few cancer types with exceptionally larger sample sizes, potentially introducing bias.

### Prevalent non-mutational inactivation of p53 in human cancers predicted using SVM models

We examined the predictive power of our SVM models by applying them to *TP53*^WT^ and *TP53*^MM^ tumors, which were excluded from the training and testing phases. To avoid potential confusion caused by the term “p53 loss of function (LoF)”, which typically refers to p53 dysfunction due to mutations, we designated samples predicted to be *TP53*^TM^-like samples as “reduced function (RF)” instead. The results revealed that the majority of *TP53*^WT^ samples (87%-99%) and almost all *TP53*^MM^ samples (98%-100%) were predicted as RF in all seven cohorts: LUNG (94%, 100%), BRCA (87%, 98%), COAD (99%, 100%), HNSC (96%, 100%), STAD (96%, 98.17%), UCEC (97%, 99%), LIHC (90%, 100%) (Supplementary Table 4). Using the LUNG cohort as an example, 94% (310 out of 330) of the *TP53*^WT^ samples and 100% (n = 419) of *TP53*^MM^ samples were predicted to be RF (**Figure 3 a)**. These results indicate the prevalence of non-mutational inactivation of p53 in human cancers.

**Figure 3.**
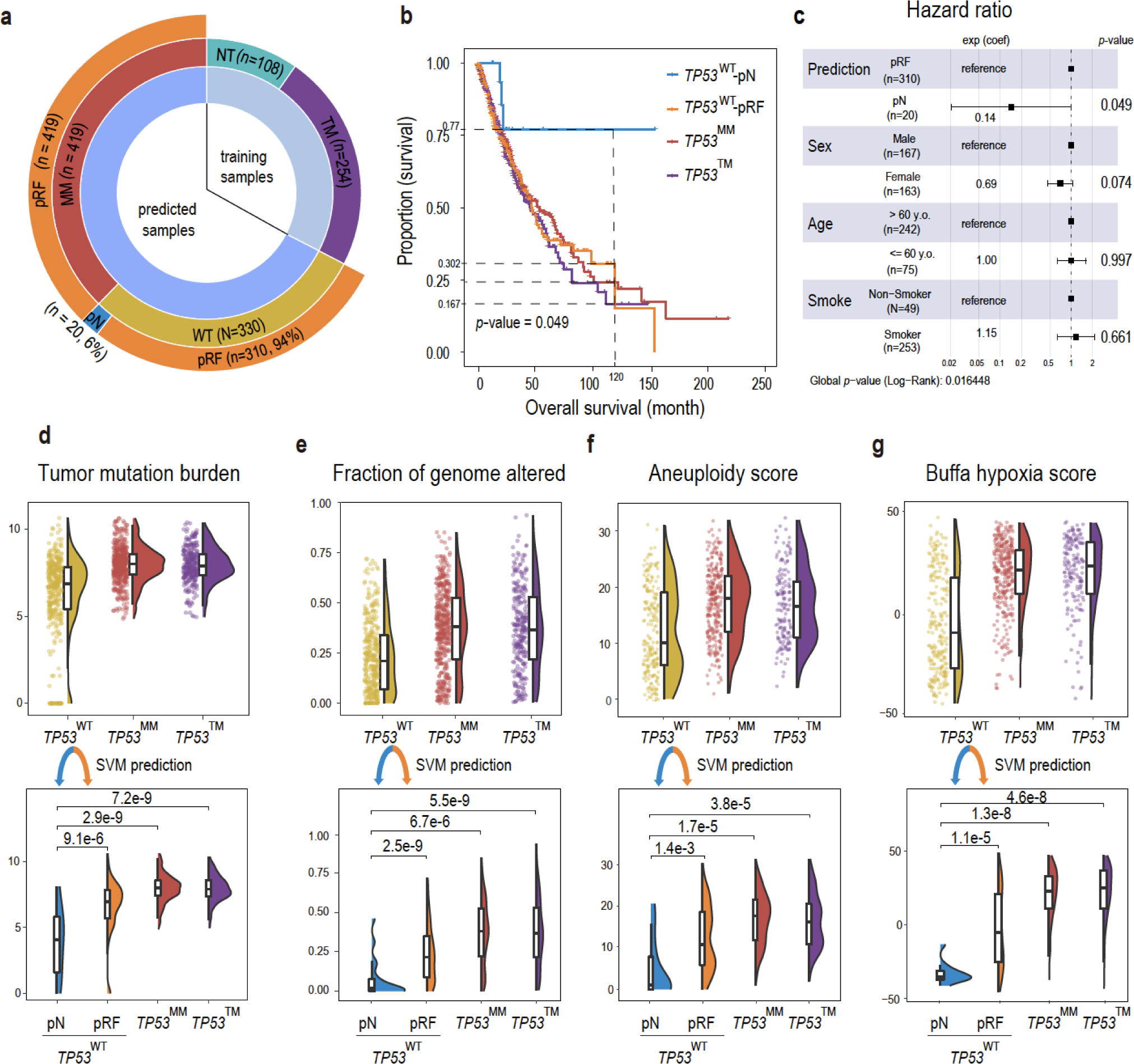
Evaluation of the SVM predictions in the TCGA lung cancer cohort. (**a**) A sunburst chart showing the breakdown of the lung cancer samples before (inner and middle layer circles) and after (outer layer circle) SVM prediction. Training samples consist of NT (adjacent normal tissue) and *TP53*^TM^ (tumor tissue with *TP53* truncating mutation) groups, representing p53-normal and p53-LoF status, respectively. Samples that were subjected to SVM prediction include *TP53*^WT^ (tumor tissue with wild-type *TP53*) and *TP53*^MM^ (tumor tissue with *TP53* missense mutation or in-frame mutation) groups. Samples in *TP53*^WT^ and *TP53*^MM^ groups were predicted as normal (pN) or RF (pRF) by the SVM model. (**b**) Overall survival of patients with *TP53*^MM^, *TP53*^TM^, *TP53*^WT^-pN (wild-type *TP53* predicted as normal) and *TP53*^WT^-pRF (wild-type *TP53* predicted as reduced function) tumors. (**c**) Forest plot of a Cox Proportional-Hazards model showing *p*-values, hazard ratios, and the 95% confidence intervals for covariates. (**d**) Upper panel: Comparison of tumor mutational burden (TMB) among *TP53*^WT^, *TP53*^MM^, and *TP53*^TM^ groups. Lower panel: the *TP53*^WT^ tumors were further divided into two subgroups (*TP53*^WT^-pN and *TP53*^WT^-pRF) based on SVM prediction. (**e**) Similar to (d), comparison of the copy number variation burden measured by “Fraction of genome altered”. (**f**) Similar to (d), comparison of the aneuploidy score. (**g**) Similar to (d), comparison of hypoxia score.

To assess the prognostic value of the SVM prediction, we analyzed the LUNG cohort by dividing *TP53*^WT^ tumors into two subgroups: *TP53*^WT^-pRF, representing *TP53*^WT^ samples predicted as “reduced p53 function”, and *TP53*^WT^-pN, representing *TP53*^WT^ samples predicted as “normal p53 function”. Our analysis revealed that the 10-year overall survival (OS) rates for *TP53*^WT^-pN (n = 20) and *TP53*^WT^-pRF (n = 310) were 77.0% and 30.2%, respectively. As reference points, the 10-year OS rates for *TP53*^MM^ (n = 419) and *TP53*^TM^ (n = 254) were 24.3% and 16.7%, respectively (**Figure 3 b**). It is noted that the difference of OS rates between lung cancer patients with *TP53*^WT^-pN tumors and those with *TP53*^WT^-pRF tumors became more significant after adjusting for demographic variables such as sex, age, and smoking status using Cox regression (*P* = 0.016, **Figure 3 c**). In contrast, the OS rate of patients with *TP53*^WT^-pRF tumors did not differ significantly from that of patients with *TP53*^MM^ or *TP53*^TM^ tumors. The TCGA breast cancers were divided into five subtypes including “Basal”, “HER2+”, “Luminal A”, “Luminal B” and “Normal-like” (Supplementary Figure 5 a). Most (71 out of 84 or 85%) of the *TP53*^WT^-pN tumors are classified as Luminal-A subtype, but we did not observe a significant difference in progression-free survival (PFS) of patients with this subtype. The remaining 15% (13 out of 84) of *TP53*^WT^-pN cases are normal-like breast tumors. Within this subgroup, the *TP53*^WT^-pRF patients exhibited a significantly reduced PFS compared to the *TP53*^WT^-pN group (*P* = 0.017, log-rank test) (Supplementary Figure 5 f). We were unable to perform comparative analyses for “Basal”, “HER2+”, and “Luminal B” subtypes as there were almost no *TP53*^WT^-pN cases. Normal-like tumors were generally considered as “artifacts” resulting from a high percentage of normal specimens or slow-growing basal-like tumors [44]. However, our data suggested that the PFS is significantly reduced when p53 function is compromised, supporting the classification of normal-like tumors as a distinct subtype rather than mere normal tissue contamination. In summary, in both lung and breast (normal-like) cancers, *TP53*^WT^-pRF tumors exhibit a significantly worse prognosis compared to the *TP53*^WT^-pN tumors.

Considering the crucial role of p53 in DNA damage repair [45, 46], we hypothesized that p53-defective tumors would accumulate more DNA damage compared to p53 normal tumors. To investigate this hypothesis, we compared measurements of genome and chromosome instability, including tumor mutational burden (TMB), copy number variation burden, and aneuploidy score, between the *TP53*^WT^-pRF and *TP53*^WT^-pN groups. The analysis revealed significantly higher levels of genome instabilities in the *TP53*^WT^-pRF group compared to the *TP53*^WT^-pN group, indicating deficient DNA damage repair in *TP53*^WT^-pRF tumors (**Figure 3 d-e**). Similar trends were observed in the other TCGA cohorts (Supplementary Table 5). It is known that p53 can decrease cell hypoxia (i.e., insufficient oxygen in the tumor microenvironment) by inhibiting HIF1A activity [47-49]. Consistent with these findings, the Buffa hypoxia score [50] was significantly lower in the *TP53*^WT^-pN tumors compared to the *TP53*^WT^-pRF, *TP53*^MM^ and *TP53*^TM^ tumors (**Figure 3 f**). Overall, *TP53*^WT^-pRF tumors exhibited increased genomic instability, worse prognosis, and higher hypoxia levels, resembling the p53 mutant tumors. These data not only functionally reaffirm our SVM predictions but also reveal the potential limitation of determining *TP53* status based solely on DNA sequencing.

### Increased sensitivity of *TP53*^WT^-pRF tumors to chemo- and radiation therapy

The impact of mutant p53 on the response to chemo- and radiation therapy remains controversial. Several studies have associated p53 mutations with reduced sensitivity to these therapies [10, 11, 51-53], while other studies have suggested that p53 inactivation actually increases the tumor’s sensitivity [12-14]. Unfortunately, direct comparisons of therapy efficacy using TCGA samples are not feasible due to the lack of treatment response data. To overcome this limitation, we assessed the chemo- and radiation sensitivity using previously published gene signatures (Supplementary Table 6) and conducted investigations with preclinical animal models to examine the therapeutic effects of radiation. Specifically, we evaluated chemotherapy sensitivity in LUNG and BRCA cohorts using RPS (recombination proficiency score) which has been clinically validated in breast cancer and NSCLC (non-small cell lung cancer) patients [54]. For the assessment of tumor radiation sensitivity, we utilized RSS (radiation sensitivity signature) score and tested in breast cancer [55] (see Methods). To explore the relationship between predicted p53 status and the response to RT (radiation therapy) *in vivo*, we analyzed an independent dataset consisting of 35 PDX (patient-derived xenografts) models that closely mimic the genetic and phenotypic characteristics of glioblastoma (GBM) patients [56].

When comparing chemo-sensitivity across groups with different p53 statuses in the LUNG cohort, we found that tumors in the NT group exhibited the highest RPS, while tumors in the *TP53*^MM^ and *TP53*^TM^ groups displayed the lowest RPS. Expectedly, the RPS scores of *TP53*^WT^-pN tumors were close to those of the NT group and were significantly higher than the *TP53*^WT^-pRF tumors (*P* = 2.8×10^-9^, two-sided Wilcoxon test). Similar results were observed in the BRCA cohort. Notably, four BRCA *TP53*^MM^ tumors that were predicted to be p53 normal also had significantly higher RPS scores than the remaining *TP53*^MM^ samples (*P* = 0.029, two-sided Wilcoxon test) (**Figure 4 a-b)**. The lower RPS score was linked to increased mutagenesis, adverse clinical features, and inferior patient survival rates, but such adverse prognosis could be counteracted by adjuvant platinum-based chemotherapy [54]. While mutations in *BRCA1*/*BRCA2* are commonly known as the primary drivers of homologous recombination deficiency (HRD), strong positive associations between the *TP53* mutation ratio and HRD scores have been observed in TCGA pan-cancer analysis [57]. Moreover, studies have indicated the involvement of p53 in regulating homologous recombination [58, 59]. These data suggested that platinum-based chemotherapy may provide greater benefits to *TP53*^WT^-pRF tumors compared to *TP53*^WT^-pN tumors.

**Figure 4.**
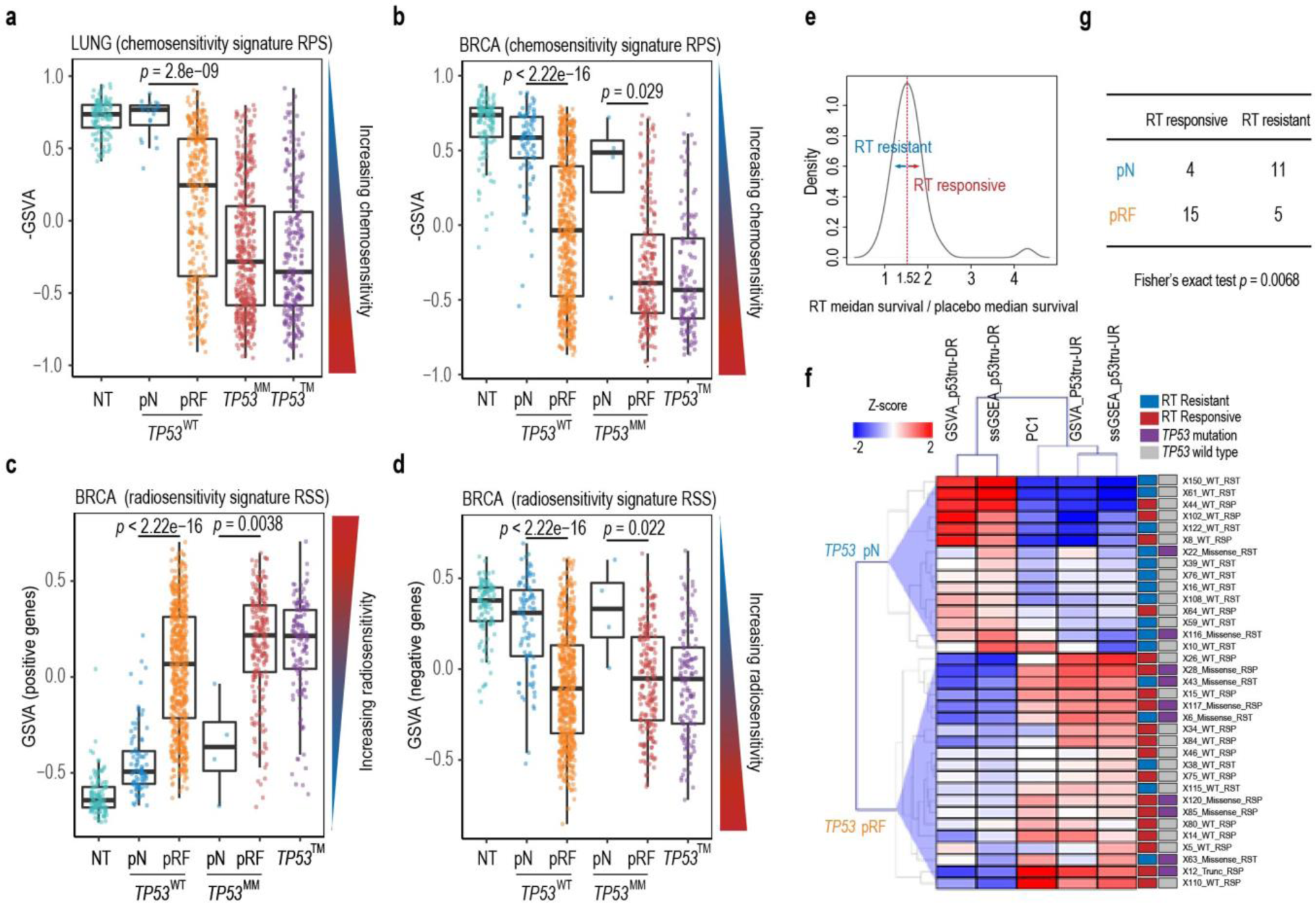
Relationships between p53 status and chemo-and radiation therapy sensitivities. (**a-b**) Comparison of the RPS scores (chemosensitivity signature, represented by negative GSVA score, see Methods) amongst the NT, *TP53*^WT^-pN, *TP53*^WT^-pRF, *TP53*^MM^, and *TP53*^TM^ groups in the TCGA LUNG and BRCA cohorts, respectively. (**c-d**) Comparison of the RSS scores (radiosensitivity signature, represented by GSVA score, see Methods) of positive and negative genes amongst NT, *TP53*^WT^-pN, *TP53*^WT^-pRF, *TP53*^MM^-pN, *TP53*^MM^-pRF, and *TP53*^TM^ groups in the BRCA cohort, respectively. (**e**) Distribution of the ratio of median survival days between the RT (radiation therapy) group and the placebo group. PDX models with ratios >= 1.52 were considered as RT-responsive. (**f**) A heatmap showing the p53 functional status predicted by unsupervised clustering using composite expression (GSVA, ssGSEA, and PC1) of GBM-specific p53tru-DR and p53tru-UR genes in PDX data. RT response and *TP53* mutation status of each sample are color coded on the right side of the heatmap. (**g**) A contingency table shows that pRF (predicted reduced function) samples are significantly associated with RT responsiveness.

We assessed radiosensitivity by dividing the RSS gene signature into positive (i.e., genes positively correlated with radiosensitivity) and negative (genes negatively correlated with radiosensitivity) subsets. Our analysis focused on the TCGA BRCA cohort since the RSS signature was derived from breast cancer. When comparing *TP53*^WT^-pN tumors with *TP53*^WT^-pRF tumors using positive genes, we found a significant increase in RSS scores for *TP53*^WT^-pRF tumors (**Figure 4 c**), indicating heightened radiosensitivity. Meanwhile, when measured using negative genes, *TP53*^WT^-pRF tumors showed significantly decreased RSS scores, also suggesting increased radiosensitivity (**Figure 4 d**). On the other hand, *TP53*^MM^-pN tumors displayed significantly reduced radiosensitivity compared to *TP53*^MM^-pRF tumors (**Figure 4 c-d**).

Although there is one gene (*RAD51*) from the RPS signature and four genes from the RSS signature overlapped with the SVM feature genes for LUNG or BRCA (Supplementary Figure 6 a), we obtained consistent results even after excluding these overlapped genes (Supplementary Figure 6 b-e). These findings collectively indicate that tumors predicted to have reduced p53 function, akin to p53 mutant tumors, show increased sensitivity to platinum-based chemotherapy and radiation therapy. This could be attributed to compromised DNA damage repair due to reduced p53 function, making tumor cells more susceptible to the effects of chemo and radiation therapies.

We further confirmed this finding *in vivo* using 35 PDX models. We first defined 17 p53tru-DR and 19 p53tru-UR genes from the TCGA GBM cohort (Supplementary Table 1). Due to limited sample sizes of normal tissues and tumors with *TP53* truncating mutations, training an SVM model to predict p53 status was unfeasible. Alternatively, we employed unsupervised clustering to predict the p53 functional status using CESs calculated from RNA-seq data of PDX mouse models (Supplementary Table 7). To determine the RT responsiveness, we calculated the ratio of median survival days between the RT group and the placebo group, and PDX models with ratios >= 1.52 were considered as RT-responsive (**Figure 4 e**, Supplementary Table 7). These results indicate a significant over-representation of RT responders within the pRF group (Fisher’s exact test, *P* = 0.0068) (**Figure 4 f and g**). Although *TP53* mutations were enriched in the pRF group (**Figure 4 f)**, the association between *TP53* genetic mutation and RT response did not reach statistical significance (Fisher’s exact test, *P* = 0.13). In summary, these findings suggested that our *in silico* predictions effectively uncovered a significant association between p53 status and RT response, which would have been overlooked if solely examining the *TP53* genetic status.

### *TP53^WT^*-pRF partially attributes to WES false-negative and MDM2/MDM4 amplification

Next, we sought to explore the potential factors and mechanisms that could account for the majority of *TP53*^WT^ tumors being predicted as RF by the SVM model. We investigated the RNA and protein expression, re-evaluated all the *TP53* missense mutations (reported from WES) using RNA-seq data, and examined the alteration status of p53 upstream regulators MDM2 and MDM4.

We did not detect significant changes in p53 protein abundance between *TP53*^WT^-pN and *TP53*^WT^-pRF tumors. *TP53* RNA expressions were even increased in the *TP53*^WT^-pRF tumors (Supplementary Figure 7), which aligns with previous research indicating that mutant p53 is associated with increased *TP53* mRNA expression [60]. These results also suggest that the impaired p53 function observed in the *TP53*^WT^-pRF group is unlikely to be attributed to the reduced *TP53* mRNA and protein levels.

We then reassessed all the *TP53* missense mutations, such as R249 and R273, in the *TP53*^WT^-pRF tumors using the RNA-seq sequence data of the LUNG and BRCA cohorts. Surprisingly, 10.3% (32 out of 310) LUNG and 10.2% (57 out of 561) BRCA tumors in the *TP53*^WT^-pRF group were indeed p53 mutants, which were supported by the substantial number of RNA-seq reads carrying the mutant alleles (Supplementary Table 8). For example, two LUAD samples (TCGA-55-6987 and TCGA-55-8621), initially classified as *TP53*^WT^ by TCGA, were re-evaluated and found to possess mutant allele fractions (MAF) of 45% (44 out of 98) and 35% (35 out of 100), respectively. Notably, these two mutations were also detectable in the WES data, albeit with much lower numbers of supporting reads (2 and 5 reads, respectively). This discrepancy explains why they were not identified by the TCGA somatic variant caller (Supplementary Figure 8). The substantial increase in MAFs observed in the RNA-seq data is probably due to the preferential expression of the mutant alleles. Interestingly, when comparing the amino acid locations of missense mutations reported from WES data, these mutations identified through RNA-seq data were significantly enriched (*P* = 1.05×10^-6^, two-sided Fisher exact test) at the p53 R249 position in both lung and breast cancers (**Figure 5 a and b**). A missense mutation at the R249 position is generally recognized as a structural mutation that destabilizes the p53 protein [61]. Furthermore, the mutant tumors rescued from the RNA-seq data exhibited similar genomic and pathophysiology characteristics as the *TP53*^MM^ and *TP53*^TM^ tumors (Supplementary Figure 9).

**Figure 5.**
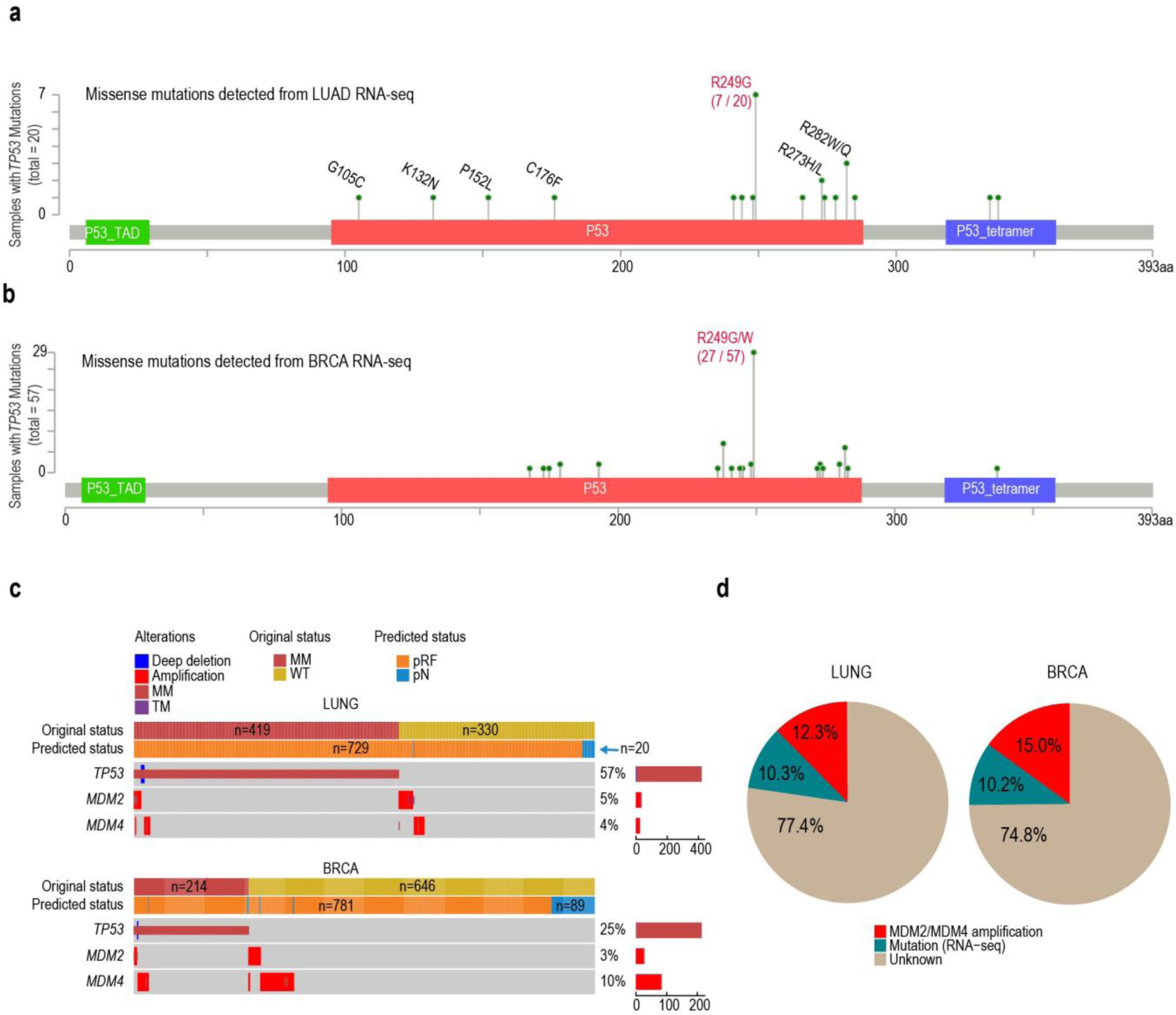
Dissection of *TP53*^WT^-pRF cases. (**a-b**) (**a-b**) Lollipop charts showing the positions of missense mutations identified using RNA-seq data but missed from the whole-exome sequencing data in TCGA lung adenocarcinoma (LUAD) and breast invasive carcinomas (BRCA), respectively. (**c**) Oncoprint plot showing the *TP53*’s original and predicted status, compared with the *MDM2* and *MDM4* amplification status. Percentages of samples with *TP53*/*MDM2*/*MDM4* alterations among the prediction set (MM and WT samples) are indicated on the right side, as well as the numbers of these samples (indicated by bar plots). (**d**) Pie charts showing the fraction of *TP53*^WT^-pRF cases that can be explained by “mutation detected from RNA-seq” (green) and “*MDM2* or *MDM4*” amplification (red).

MDM2 and its homolog MDM4 are well-documented negative regulators of p53 [62-64]. In our analysis, we found that amplification of *MDM2* and/or *MDM4* was mutually exclusive with *TP53* mutations in lung cancer (odds ratio = 4.32, *P* = 2.2×10^-16^, χ^2^ test) and breast cancer (odds ratio = 1.46, *P* = 2.0×10^-16^, χ^2^ test) (**Figure 5 c**). Consistent with this observation, all samples displaying amplification of *MDM2* and/or *MDM4* (n = 38 in LUNG, n = 84 in BRCA) were classified as pRF. These samples accounted for 12.3% and 15.0% of all *TP53*^WT^-pRF samples in the LUNG and BRCA cohorts, respectively (**Figure 5 d**). Collectively, 22-25% of *TP53*^WT^ tumors predicted as *TP53*^WT^-pRF can be attributed to false-negative results of WES assay or amplification of *MDM2* and *MDM4*.

## Discussion

Tumor suppressor genes such as *RB1*, *PTEN*, and *CDKN2A* are generally inactivated through homozygous deletions, resulting in minimal or complete loss of protein expression (https://www.cbioportal.org/). In contrast, *TP53*, unlike other tumor suppressors, predominantly undergoes missense mutations that give rise to altered or dysfunctional protein variants (https://www.cbioportal.org/). In addition, the oncogenic role of mutant p53 has been extensively characterized through multiple lines of evidence [65], including the over-expression in p53 mutants, accumulation of mutant p53 protein in the nucleus and the cytoplasm, and functional studies *in vivo* and *in vitro* systems [66-69]. The primary objective of this study is to investigate the loss or reduction of p53’s tumor suppressor function in tumors that may appear functionally normal due to the absence of *TP53* genetic alterations. Based on the assumption that p53’s tumor suppressor function remains normal in NT samples, while it is compromised or significantly reduced in TM samples, we trained SVM models using normal tissues (NT samples) and tumor samples with truncating *TP53* mutations (TM samples) that result in shortened protein. However, it is essential to recognize that our training dataset is not without imperfections, given that the p53 statuses of these samples are inferred rather than definitively determined. To be more precise, the predicted outcomes should be termed NT-like or TM-like. Looking ahead, datasets that encompass paired RNA- and DNA-sequencing information, coupled with clearly defined p53 statuses are needed to overcome the existing limitations and gain more conclusive insights. Meanwhile, *TP53* missense mutations that were predicted to be functionally reduced (*TP53*^MM^-pRF) should be interpreted as the “reduced tumor suppressor function of p53”. Nevertheless, we could not rule out the possibility that the mutant p53 protein in the *TP53*^MM^-pRF samples might gain oncogenic functions. Additionally, although p53tru-UR genes are negatively regulated by p53 through indirect and unclear mechanism, we retained these genes for downstream analyses since they can effectively predict p53’s functional status as p53tru-DR genes do. Our strategy allows us to identify tumors with compromised p53 function resulting from multiple mechanisms, and it is plausible that some cases predicted to have reduced p53 function may genuinely represent a loss of function, while others may indicate dysregulation.

Although the expression of p53-regulated genes could be potentially influenced by other factors, our *TP53*^WT^-pRF predictions, as well as the p53-regulated genes defined in this study, are highly specific to *TP53*. We performed the Fisher’s exact test on the prediction results of the LUNG and BRCA cohorts, to assess whether the mutation status (WT or mutant) of other cancer-related genes correlates with the prediction results (pN and pRF) of *TP53*^WT^ and *TP53*^MM^ samples. Notably, in LUNG, *TP53* emerged as the most significant gene associated with our prediction, with a considerably lower *p*-value compared to the other three genes meeting the threshold of *p* < 0.05. Similarly, in BRCA, *TP53* ranked as the 2^nd^ most significant gene after *CDH1* (both *TP53* and *CDH1* show considerably lower *p*-values compared to the other five genes), indicating a potential synergistic effect between *CDH1* and *TP53* and emphasizing the importance of *TP53* in *TP53*^WT^-pRF prediction. Previous studies did report the somatic co-inactivation and collaboration of *TP53* and *CDH1* in breast and other cancers [70, 71]. Additionally, p53tru-UR and -DR genes are highly specific to the p53 pathway, as evidenced by their significant enrichment in this pathway compared to others; the “p53 signaling pathway” was ranked as the most significantly enriched pathway (adjusted *p* = 1.26E-31).

According to our analyses, approximately 22-25% of *TP53*^WT^ tumors predicted as *TP53*^WT^-pRF can be attributed to false-negative results (i.e., *TP53* mutations that failed to be detected from the WES assay) or amplification of *MDM2* and *MDM4*. However, the underlying mechanisms for the remaining 75-78% of *TP53*^WT^-pRF are unknown and warrant further investigation. In order to gain more insights, we analyzed the DNA methylation data generated from the Infinium HumanMethylation450 BeadChip array. However, we did not observe significant differences in the DNA methylation patterns among all nine CpG sites within the *TP53* gene (cg02087342, cg06365412, cg10792831, cg12041075, cg12041429, cg13468400, cg16397722, cg18198734, cg22949073) when comparing *TP53*^WT^-pRF and *TP53*^WT^-pN tumors. This suggests that p53 function may be compromised through other non-mutational mechanisms such as post-translational modification [25-29], chromatin states [31], and interaction with microRNAs [32, 33]. Unfortunately, due to the unavailability of matched data, we were unable to perform specific analyses in these areas. We also investigated the potential contributions of “p53-regulated genes” and “p53 functional partners” in LUNG cohort, and found up to 54.8% of *TP53*^WT^-pRF samples (which cannot be explained by “undetected *TP53* mutation” or “*MDM2/4* amplification”) have mutations in these genes. However, the mutation frequencies of individual genes are low, and we are unable to assess the statistical significance of their associations with *TP53*^WT^-pRF.

It is noted that our data suggested that platinum-based chemotherapy, rather than all kinds of chemotherapies, may provide greater benefits to *TP53*^WT^-pRF tumors compared to *TP53*^WT^-pN tumors. Using drug imputation data of TCGA patients [71, 72], we further examined the treatment response of *TP53*^WT^-pRF samples. In the first study [71], we identified 22 drugs exhibiting elevated sensitivity (Wilcoxon rank sum test, adjusted *p*-value < 0.05) in *TP53*^WT^-pRF tumors and 5 drugs showing increased sensitivity (adjusted *p*-value < 0.05) in *TP53*^WT^-pN tumors. In the second study [72], CCLE S-model prediction identified 13 drugs with increased sensitivities in *TP53*^WT^-pN tumors; GDSC A-model prediction identified 27 drugs showing increased sensitivity in *TP53*^WT^-pN tumors and 20 drugs exhibiting elevated sensitivity in *TP53*^WT^-pRF tumors. However, there is limited overlap and poor consistency between the two studies. Therefore, chemotherapies should be dissected in detail to inform clinical therapy strategies.

In our analysis, we found no significant differences in tumor purities (measured by IHC [73]) between *TP53*^WT^-pN and other tumor samples in the LUNG cohort (*P* = 0.460, Wilcoxon rank sum test) and the BRCA cohort (*P* = 0.066, Wilcox rank sum test). This suggests that the *TP53*^WT^-pN tumors are not associated with lower tumor cellularity. Additionally, we identified four *TP53*^MM^ samples from the BRCA cohort (TCGA-E2-A1B1, TCGA-LD-A74U, TCGA-BH-A1FE, and TCGA-EW-A1P1) that were predicted to have normal p53 function (Supplementary Table 4 c). We confirmed all reported mutations in these four samples using WES and RNA-seq data and ruled out the possibilities of false positives or cross-sample contaminations. The reasons behind these findings remain unclear, and the limited sample size prevents extensive analysis.

While our analytic procedure is robust and can be applied to other datasets, building an SMV model from scratch requires a large number of normal samples and tumor samples with *TP53* truncating mutations as training data. When obtaining sufficient training data is not practical, alternative unsupervised machine-learning approaches such as K-means clustering, semi-supervised learning, principal component analysis (PCA), and neural networks could be considered.

Our analyses have revealed an intriguing aspect: the majority of *TP53*^WT^ tumors are genetically intact but exhibit functional deficiencies in p53. This discovery highlights an inherent limitation in relying solely on DNA markers for patient stratification and segmentation. It emphasizes the need for the development of “ensemble” approaches that incorporate multi-omics data to capture the full complexity of p53 functionality. By considering multiple layers of molecular information, we can gain a more comprehensive understanding of p53 status and its implications in cancer. Such integrated approaches hold great promise for enhancing our ability to characterize tumors and accurately guide personalized treatment strategies.

## Conclusion

In this study, we employed a novel approach to measure p53 activity by defining p53-regulated genes and calculating their CESs as a surrogate. By training and cross-validating SVM models using CES data from the NT (p53-normal) and TM groups (p53-RF), we demonstrated the accuracy and effectiveness of our *in silico* approach. Our comprehensive analysis revealed the prevalence of non-mutational p53 inactivation in human malignancies. Moreover, our analyses unveiled that the predicted *TP53*^WT^-pRF tumors exhibited a comparable level of genomic instability to those harboring genetic *TP53* mutation. This included a significant increase in the number of mutations, copy number alterations, and aneuploidy. Importantly, patients with *TP53*^WT^-pRF tumors showed considerably worse overall survival rates when compared to those with *TP53*^WT^-pN tumors, highlighting the prognostic value of our prediction. Furthermore, when evaluated using clinically validated signatures, *TP53*^WT^-pRF tumors demonstrated significantly heightened sensitivity to platinum-based chemotherapy and radiation therapy. This observation was verified in our preclinical animal models of GBM. Additionally, we explored potential factors contributing to the *TP53*^WT^-pRF classification, such as false-negative mutation calls and amplifications of *MDM2* and *MDM4*.

## Materials and Methods

### Data collection

TCGA somatic mutation data, and the pre-calculated fraction of genome altered (FGA) score, mutation count, aneuploidy score, and Buffa hypoxia score were downloaded from cBioPortal (https://www.cbioportal.org/). TCGA WES and RNA-seq BAM files for BRCA and LUNG were downloaded from the GDC (https://portal.gdc.cancer.gov/). TCGA level-3 RNA-seq expression data, demographic and survival data were downloaded from the Xena web server (https://xena.ucsc.edu/). Pre-calculated raw RNA-seq read counts were used to identify differentially expressed genes between the NT and TM groups. The log2-transformed FPKM (i.e., Fragments Per Kilobase of exon per Million mapped fragments) was used to calculate CESs. TPM (i.e., Transcript Per Million) was used to calculate GSVA score to evaluate chemo-and radiotherapy sensitivities. ChIP-seq data (p53 binding peaks) were downloaded from the ReMap database (https://remap.univ-amu.fr/). Gene expression data (TPM) of normal tissues were downloaded from the GTEx (release V8) data portal (https://gtexportal.org/home/).

### Identification of p53-regulated genes across different cancer types

We fist compiled a set of 147 genes that have been experimentally validated as being regulated by p53. This set includes the 116 genes identified as directly activated targets of p53 (refer to Table 1 in [8]) and 31 genes that are repressed by p53 by indirect regulation (refer to Table 2 in [8]). We then analyzed p53 ChIP-seq data from ReMap2020 [38] to verify p53 binding within the gene body or the promotor region. The basal level expression of p53 regulated genes in normal tissues was evaluated using the RNA-seq data of GTEx [74]. Only genes with median TPM > 1 were included in our downstream analyses. To identify feature genes whose expression could reflect p53’s functional status in each cancer type, we performed differential expression analysis for the 147 genes in each cancer by comparing the NT group (normal tissue samples) to the TM (samples harboring *TP53* truncating mutations) using DESeq2 [75]. The significance threshold was set at an adjusted *p*-value ≤ 0.05 and |fold change| ≥ 2. We defined “p53tru-DR” genes as those showing downregulation in p53-truncated samples compared to p53-normal samples. Conversely, “p53tru-UR” genes referred to genes showing upregulation in p53-truncated samples. To verify our gene set, we included an independent LUAD dataset of East Asians (https://src.gisapps.org/OncoSG_public/study/summary?id=GIS031) into our analysis [76]. This dataset includes 21 p53-truncated samples and 88 adjacent normal lung tissues. Employing the same procedure, we identified differentially expressed genes out of 147 p53 target candidates from this data set, and compared the overlap of p53-regulated genes between this dataset and the TCGA LUNG cohort. Additionally, using the CESs calculated from p53-regulated genes identified in TCGA LUNG cohort, we predicted the p53’s functional status in this independent dataset by unsupervised clustering.

### Computation of composite expression scores

We employed four algorithms to calculate the CES, namely: (1) Gene Set Variation Analysis (GSVA) [34]; (2) single-sample GSEA (ssGSEA) [35]; (3) The first Principal Component (PC1) score of the principal component analysis (PCA); and (4) combined Z-score [36, 37]. To calculate the combined Z-score, gene expression values (log2 (FPKM)) of each sample were converted into Z-scores by Z = (x – μ)/σ, where μ and σ is the average and standard deviation of log2 (FPKM) across all samples of a gene. Given a gene set γ = {1,…,k} with standardized values *z_1_*,…,*z_k_*, for each gene in a specific sample, the combined Z-score Z_γ_ for the gene set γ is defined as:

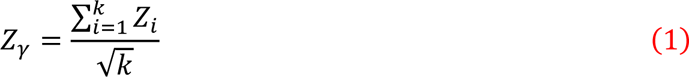

GSVA, ssGSEA, and Z-score were calculated separately for p53tru-DR and p53tru-UR genes, while PC1 was computed using all p53-regulated genes, which resulted in a total of seven CESs for each sample. The GSVA package (https://www.bioconductor.org/packages/release/bioc/html/GSVA.html) was used to calculate GSVA, ssGSEA, and Z-score. The Python package sklearn (https://scikit-learn.org/stable/) was used to perform PCA analyses.

### Training and evaluating the performance of SVM models

SVM models with linear kernel were trained using the CESs of NT (normal tissue, coded as “0”) and TM (truncating mutations, coded as “1”) groups. GridSearch was employed to pick the proper C and gamma parameters for the SVM model. We used TCGA data for both training and testing; no separate or external validation set was used. Specifically, the number of training and testing samples (n) were 362, 246, 99, 206, 104, 87, 91 and 1335 for the LUNG, BRCA, COAD, HNSC, STAD, UCEC, LIHC and pan-cancer cohorts, respectively (Supplementary Table 4 a). A total of seven features (GSVA, ssGSEA and Z-score for p53tru-DR and p53tru-UR genes respectively, and PC1 for all feature genes) were used in the SVM models. Therefore, *p* << *n* for all SVM models built in this study, where *p* represents the number of features (predictor variables) and *n* represents the number of samples.

We evaluated the performance of the SVM model of individual cancer types as the following steps. Firstly, we employed the “train_test_split” function from the “sklearn.model_selection” class to partition the samples—comprising both NT and TM samples—into a training set (constituting 75% of the data) and a separate testing set (comprising the remaining 25%). Subsequently, we performed the differential analyses to select feature genes (p53-regulated genes) with samples from the training set. This approach ensures that the selected feature genes are based solely on the training data, providing more convincing and independent evaluation results. Finally, we built the SVM models and used the testing set to evaluate the performance. We repeated this process ten times to mitigate the potential variability in the outcomes of cross-validation. In each iteration, a confusion matrix was made, and performance measurements (i.e., sensitivity/recall, precision, accuracy) were calculated. We summarized the performance of models with the mean of the measurement scores and the ROC (receiver operating characteristic) curves. The performance measurements are defined as below:

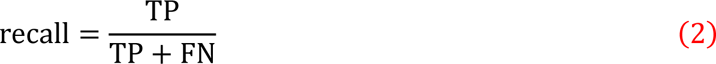

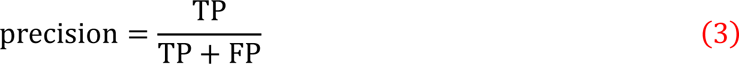

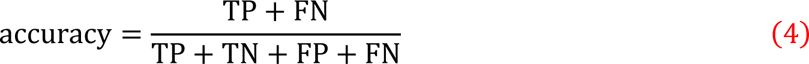

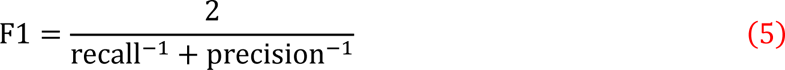

Where FN = false negative; FP = false positive; TN = true negative; TP = true positive. We used the Scikit-learn (www.scikit-learn.org) Python package for SVM modeling, ten-time hold-out validation. Details of SVM models are available in Supplementary Table 9. All TCGA sample barcodes for training and prediction used in this study are listed in Supplementary Table 4 b-f. Python source code is available from https://github.com/liguowang/epage.

### Re-evaluation of *TP53* mutation status using RNA-seq

The genomic positions of *TP53* mutations in LUNG and BRCA samples were downloaded from the cBioPortal (https://www.cbioportal.org/). The reference and mutant allele counts were then calculated from the RNA-seq BAM file using Samtools’[77]. Samples with fewer than 10 mapped reads at a given genome site were excluded, and a cut-off of minor allele frequency = 0.1 was employed to determine the genotype. A sample was considered *TP53* mutated if it had at least five high-quality (Phred-scaled sequencing quality and mapping quality > 30) reads supporting the mutant allele. The Integrative Genomics Viewer (IGV) was used to manually inspect the variants.

### Investigation of two independent treatment-related signatures

Tumor chemotherapy sensitivity (i.e., RPS score) was estimated based on 4 genes involved in DNA repair (*RIF1*, *PARI/F2R*, *RAD51*, and Ku80/*XRCC5*) reported by Pitroda *et al.* [54]. Tumor radiation sensitivity signature (RSS) scores in breast cancer were calculated based on 51-gene panel reported by Speers *et al.* [55], and the genes were divided into “positive” and “negative” groups according to the correlation between their expression level and radiation sensitivity. The R package GSVA [34] was used to calculate the RPS and RSS scores. Notably, according to the study of Pitroda *et al.*, the original RPS was defined as the sum of the expression levels times -1 after log2 transformation and Robust Multi-array Average (RMA)-normalization. In our study, RPS was represented by negative GSVA score calculated from TPM.

## GBM derived xenografts

Glioblastoma patient-derived xenografts (PDX) were generated by the Mayo PDX National Resource [56]. Mice with established orthotopic tumors were randomized into groups of 5–10 mice and treated with placebo and radiation therapy (RT). The raw sequencing data were available from the NCBI Sequence Read Archive with accession numbers PRJNA543854 and PRJNA548556. Annotated genomic and transcriptomic data are also publicly accessible through the cBioPortal (https://www.cbioportal.org/study/summary?id=gbm_mayo_pdx_sarkaria_2019). To determine the RT responsiveness of PDX, we calculated the ratio of median survivals between the RT group and placebo group. A cutoff value (1.52) was determined as the changing point on the non-parametric kernel density curve (Supplementary Table 7).

## Code Availability

Python source code of our p53 status prediction method is available from https://github.com/liguowang/epage.

## CRediT author statement

Qianpeng Li: Data curation, Formal analysis, Investigation, Methodology, Visualization, Writing - original draft, Writing - review & editing; Yang Zhang: Data curation, Formal analysis, Investigation, Methodology, Software, Visualization, Writing - review & editing; Sicheng Luo: Formal analysis, Writing - review & editing; Zhang Zhang: Writing - review & editing; Ann L. Oberg: Writing - review & editing; David E. Kozono: Writing - review & editing; Hua Lu: Writing - review & editing; Jann N. Sarkaria: Writing - review & editing; Lina Ma: Conceptualization, Project administration, Writing - review & editing; Liguo Wang: Conceptualization, Project administration, Writing - review & editing. All authors have read and approved the final manuscript.

## Competing interests

The authors declare no competing interests.

## Acknowledgments

This work is partially supported by the US National Institute of Health [U10-CA180882-07], the Center for Individualized Medicine of Mayo Clinic, the Strategic Priority Research Program of the Chinese Academy of Sciences [XDB38030400], and the Youth Innovation Promotion Association of Chinese Academy of Sciences [2019104]. The results shown in this study are in part based upon data generated by the TCGA Research Network (https://www.cancer.gov/tcga). We thank TCGA’s specimen donors and research groups that make genomic data publicly available.

## Supplementary figures

**Supplementary Figure 1.**
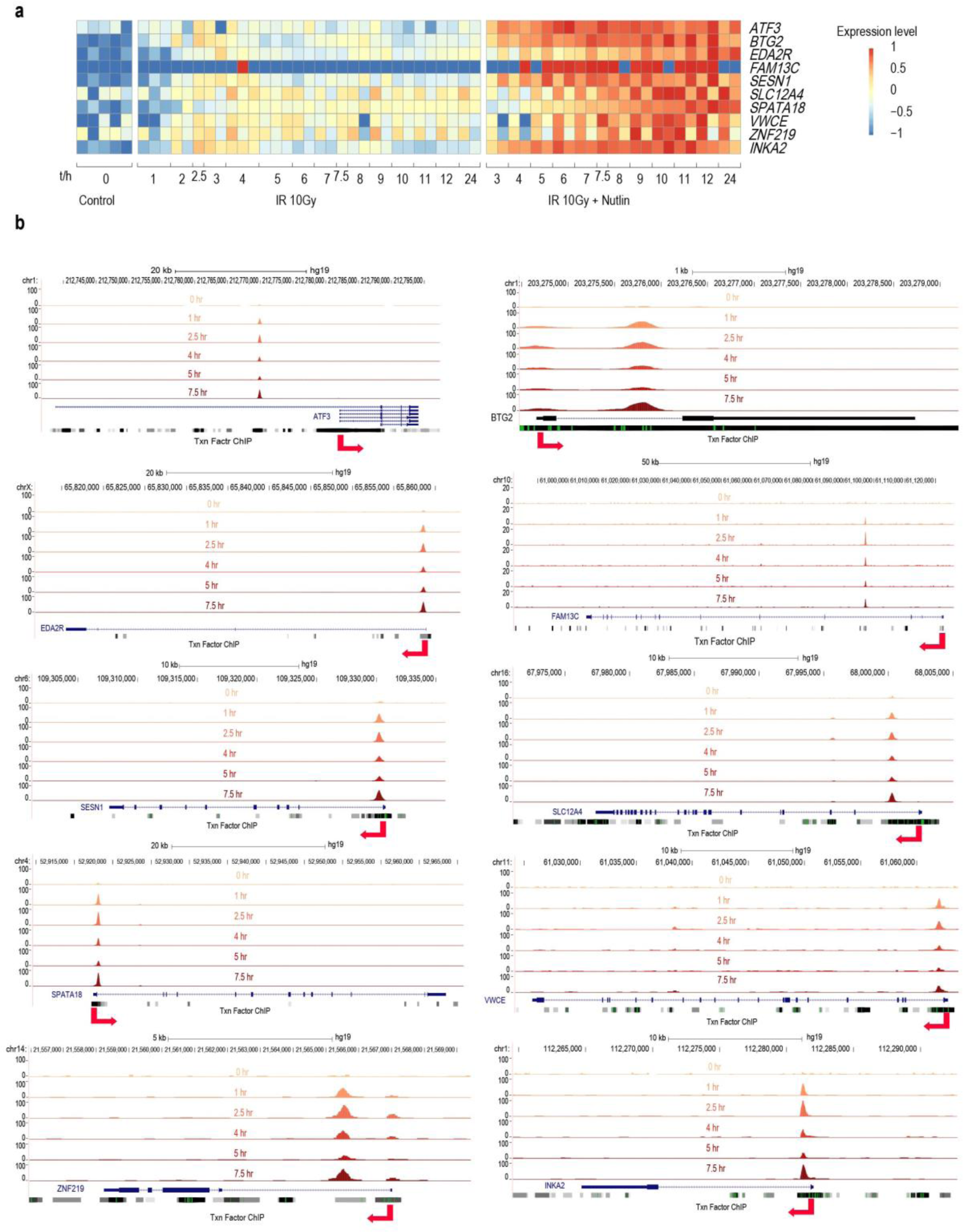
Validation of p53 target genes in γ irradiated MCF-7 cells harboring wild-type *TP53*. (**a**) Changes of expression levels of p53 targets after γ irradiation, and γ irradiation with nutlin treatment. (**b**) Visualization of p53 binding sites (p53 ChIP-seq signals) around the promoter regions of genes in (**a**).

**Supplementary Figure 2.**
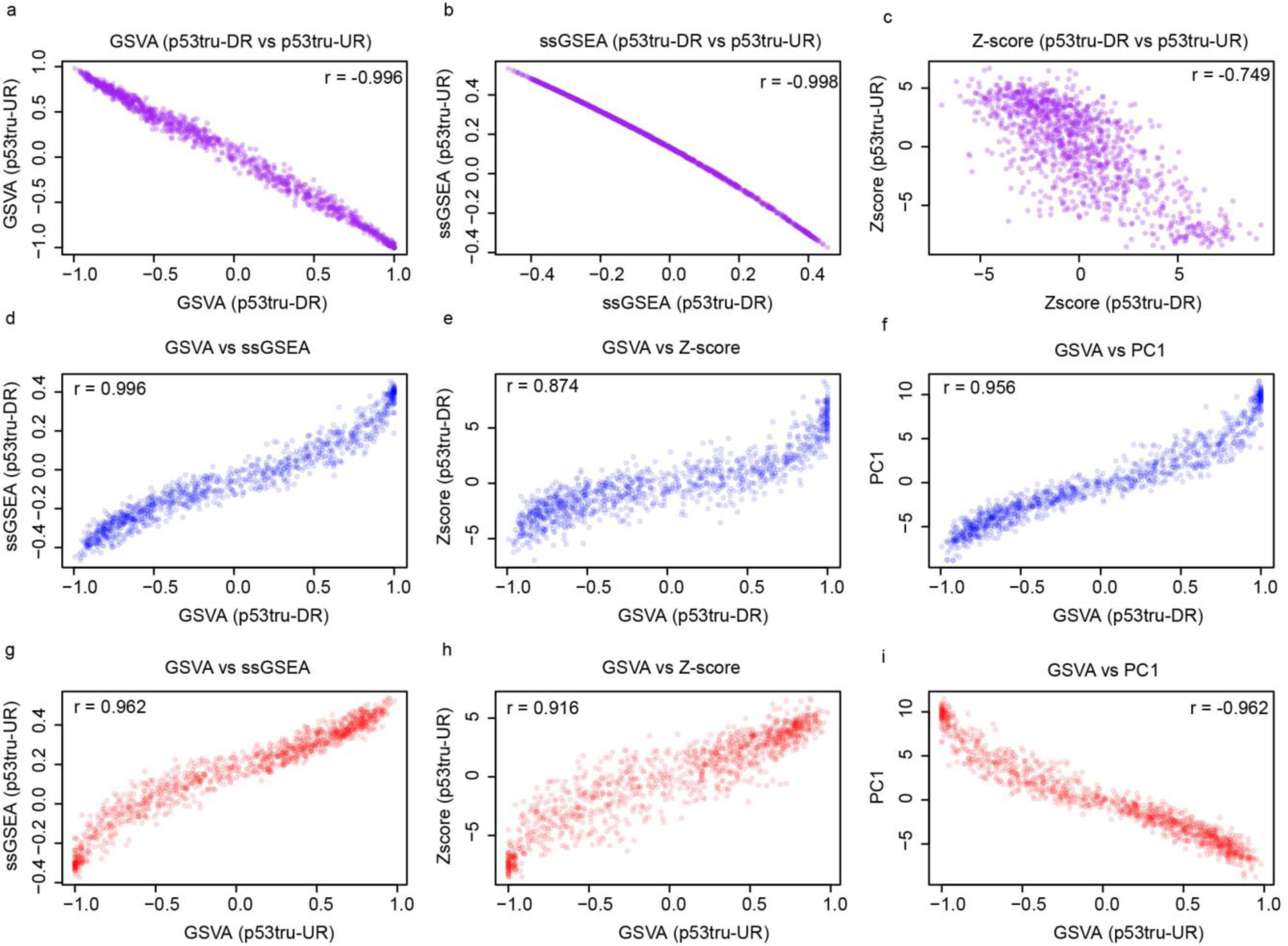
Correlations between different CESs. (**a**-**c**) Significant inverse correlations between CESs calculated from p53tru-DR and p53tru-UR genes for GSVA, ssGSEA, and Z-score. Significant positive correlations between different CESs for p53tru-DR genes (**d**-**f**) and between that for p53tru-UR genes (**g**-**h**). (**i**) Significant inverse correlation between PC1 and the GSVA of p53tru-UR genes.

**Supplementary Figure 3.**
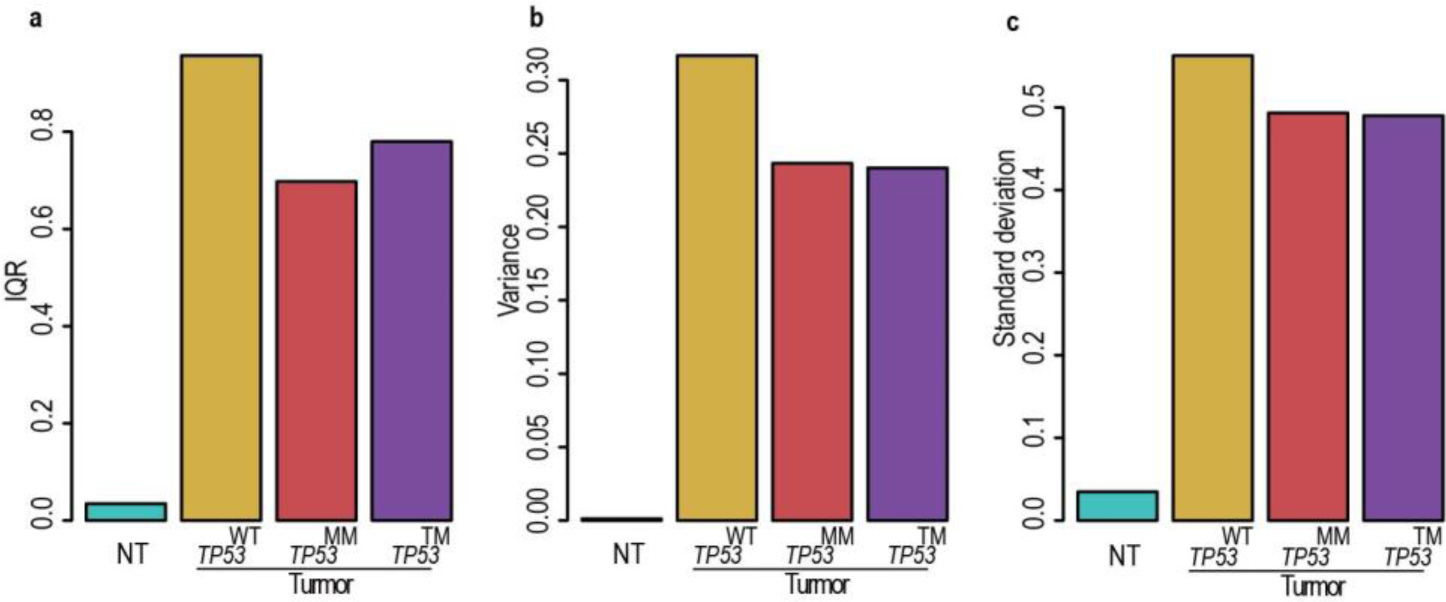
Comparison of CES distributions among different *TP53* groups in lung cancer. Bar plots show the Interquartile Range (**a**), Variance (**b**), and Standard Deviation (**c**) of CESs in NT, WT, MM, and TM groups.

**Supplementary Figure 4.**
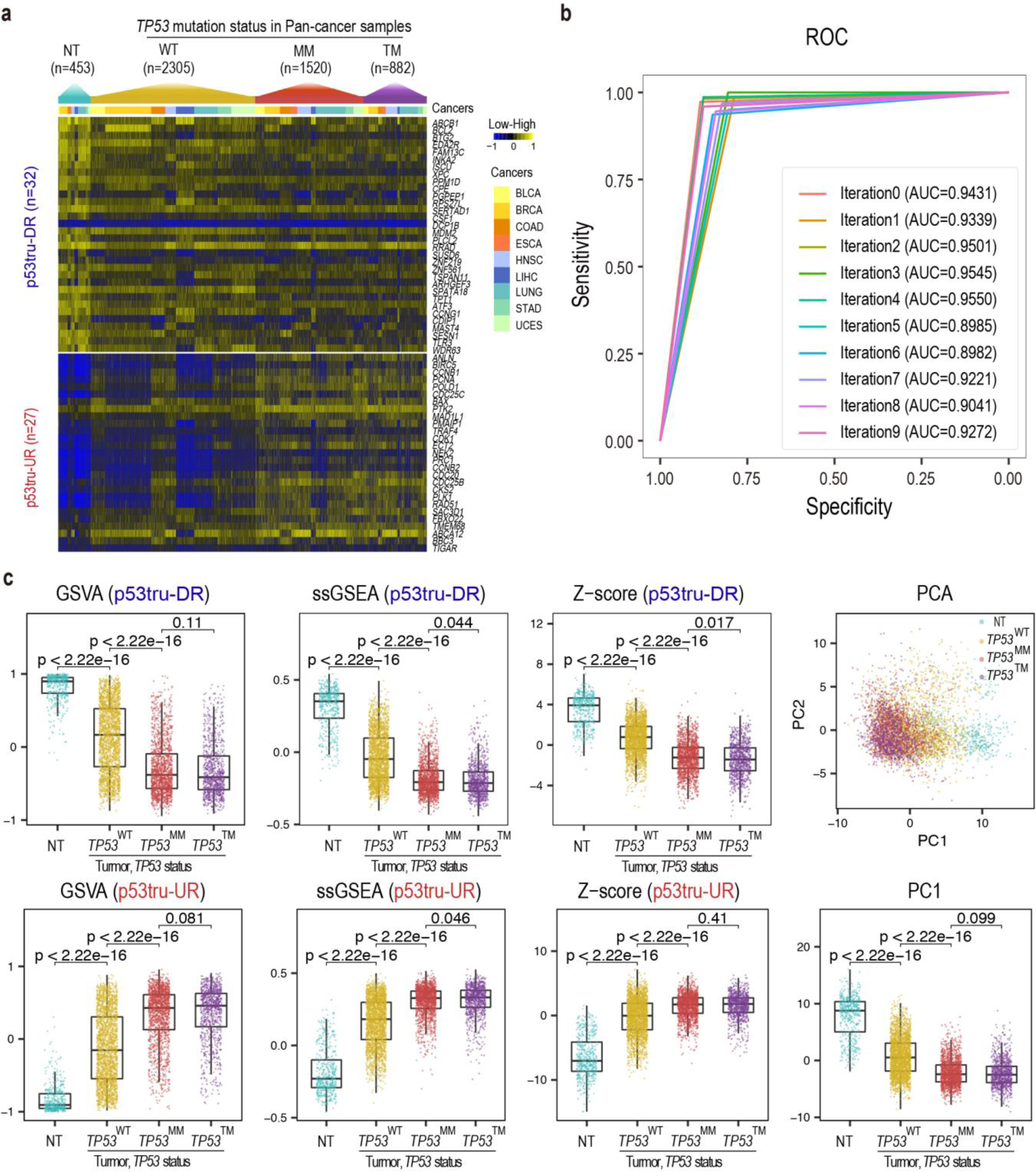
Pan-cancer analysis. (**a**) Heatmap shows the expression profiles of p53tru-DR genes and p53tru-UR genes across 9 cancer types. (**b**) ROC curve of the SVM model built from the pan-cancer cohort. (**c**) CESs calculated from p53tru-DR genes and p53tru-UR genes by different algorithms in pan-cancer cohort (similar to Figure 2).

**Supplementary Figure 5.**
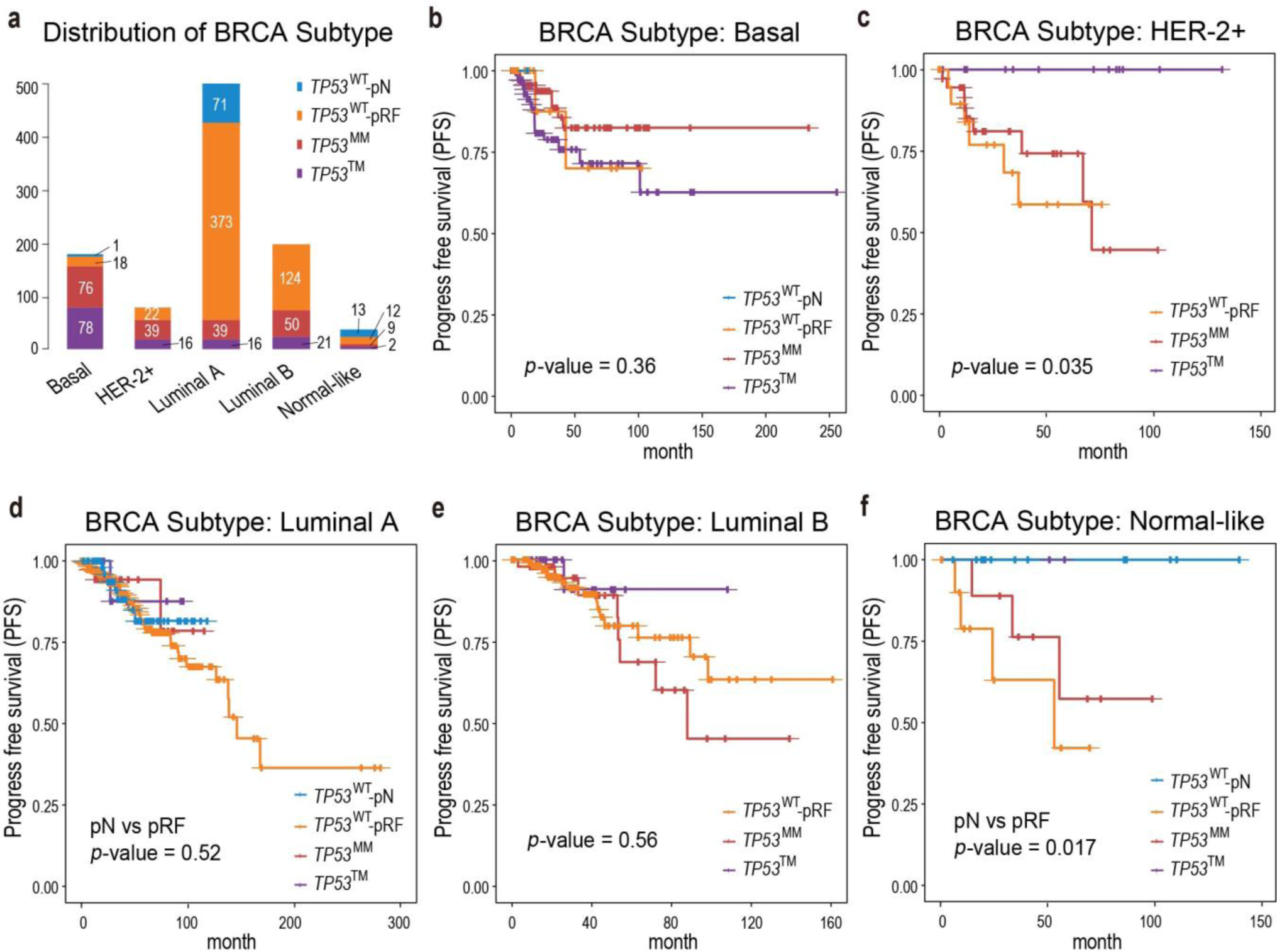
Progression-free survival (PFS) comparison among *TP53*^MM^, *TP53*^TM^, *TP53*^WT^-pN and *TP53*^WT^-pRF patients of TCGA BRCA cohort. (**a**) Bar plot shows the breakdown of BRCA patients into five subtypes, including “Basal/triple negative”, “HER2^+^”, “Luminal A”, “Luminal B” and “Normal-like”. Each subtype is further divided into four subgroups based on *TP53* status. (**b-f**) Comparison of PFS for Basal, HER2^+^, Luminal A, Luminal B, and Normal-like subtype, respectively.

**Supplementary Figure 6.**
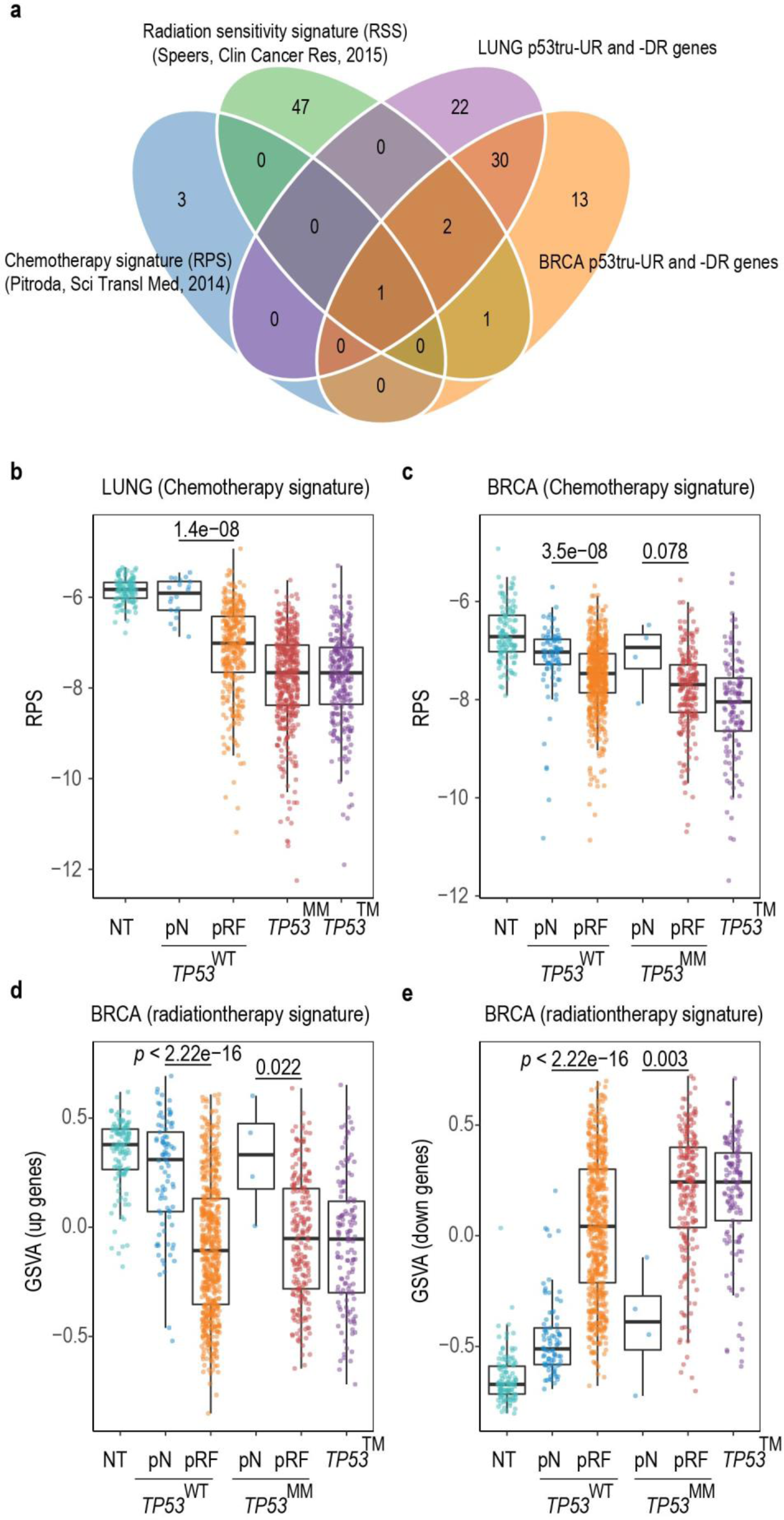
Relationships between p53 status and chemo-and radiation therapy sensitivities after removing the overlapped genes. (**a**) Venn diagram shows the overlap between p53tru-UR and p53tru-DR genes used in the LUNG and the BRCA SVM models, the chemotherapy gene signature (recombination proficiency score, RPS), and the radiation sensitivity signature (RSS). (**b**-**c**) Comparison of the RPS scores (represented by -GSVA here, see Methods) amongst NT, *TP53*^WT^-pN, *TP53*^WT^-pRF, *TP53*^MM^, and *TP53*^TM^ in the TCGA LUNG and BRCA cohort, respectively (similar to Figure 4 a-b). (**d**-**e**) Comparison of the RSS scores amongst NT, *TP53*^WT^-pN, *TP53*^WT^-pRF, *TP53*^MM^-pN, *TP53*^MM^-pRF and *TP53*^TM^ in the TCGA BRCA cohort (similar to Figure 4 c-d). GSVA was used to calculate the PRS and RSS scores. Genes overlapped with p53tru-DR or p53tru-UR genes of either LUNG or BRCA were removed when calculating the RPS and RSS scores.

**Supplementary Figure 7.**
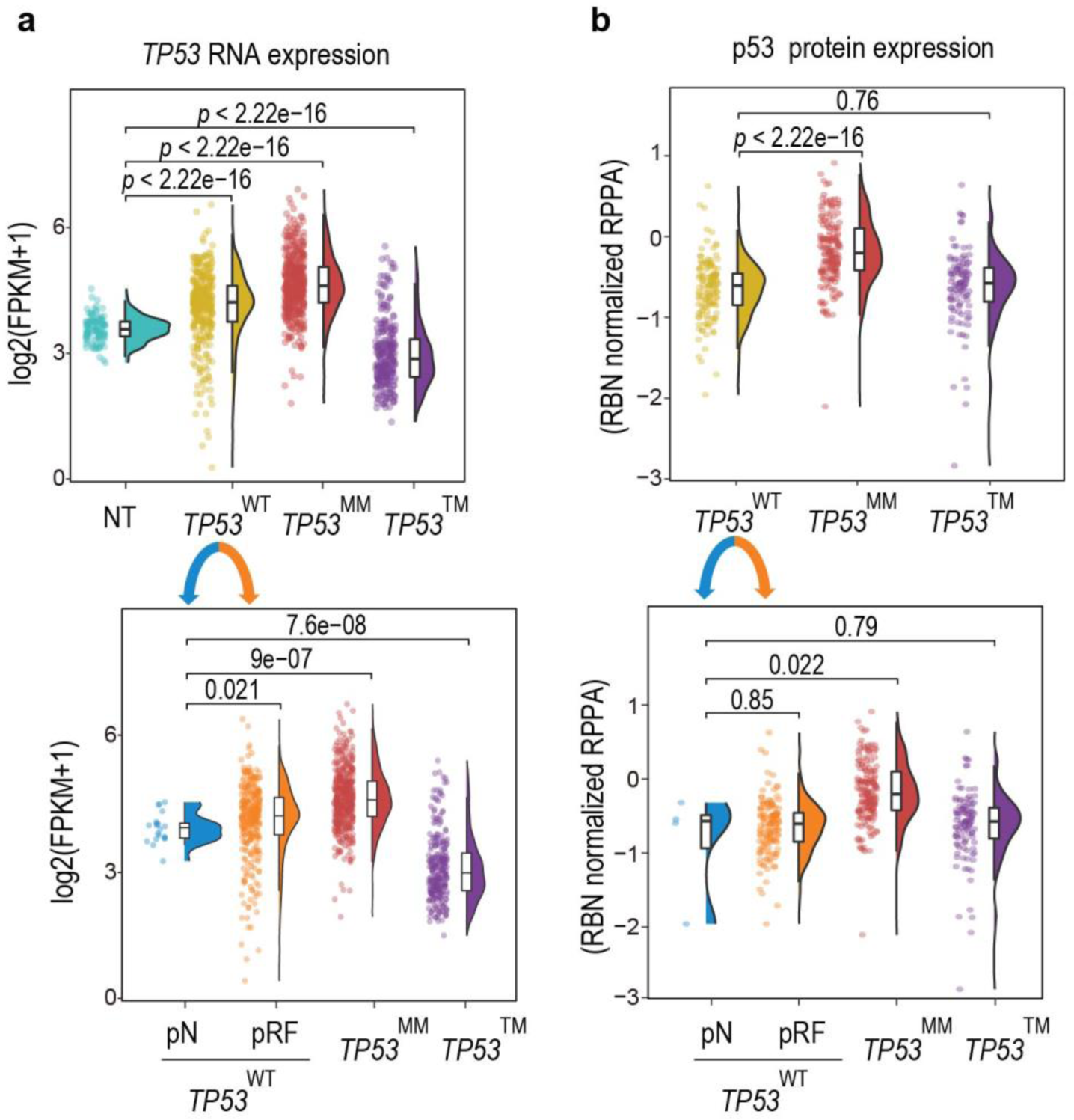
Comparison of p53 expression levels between *TP53*^WT^-pRF and *TP53*^WT^-pN samples in LUNG cohort. (**a**) Comparison of the *TP53* RNA expression. (**b**) Comparison of the p53 protein abundance.

**Supplementary Figure 8.**
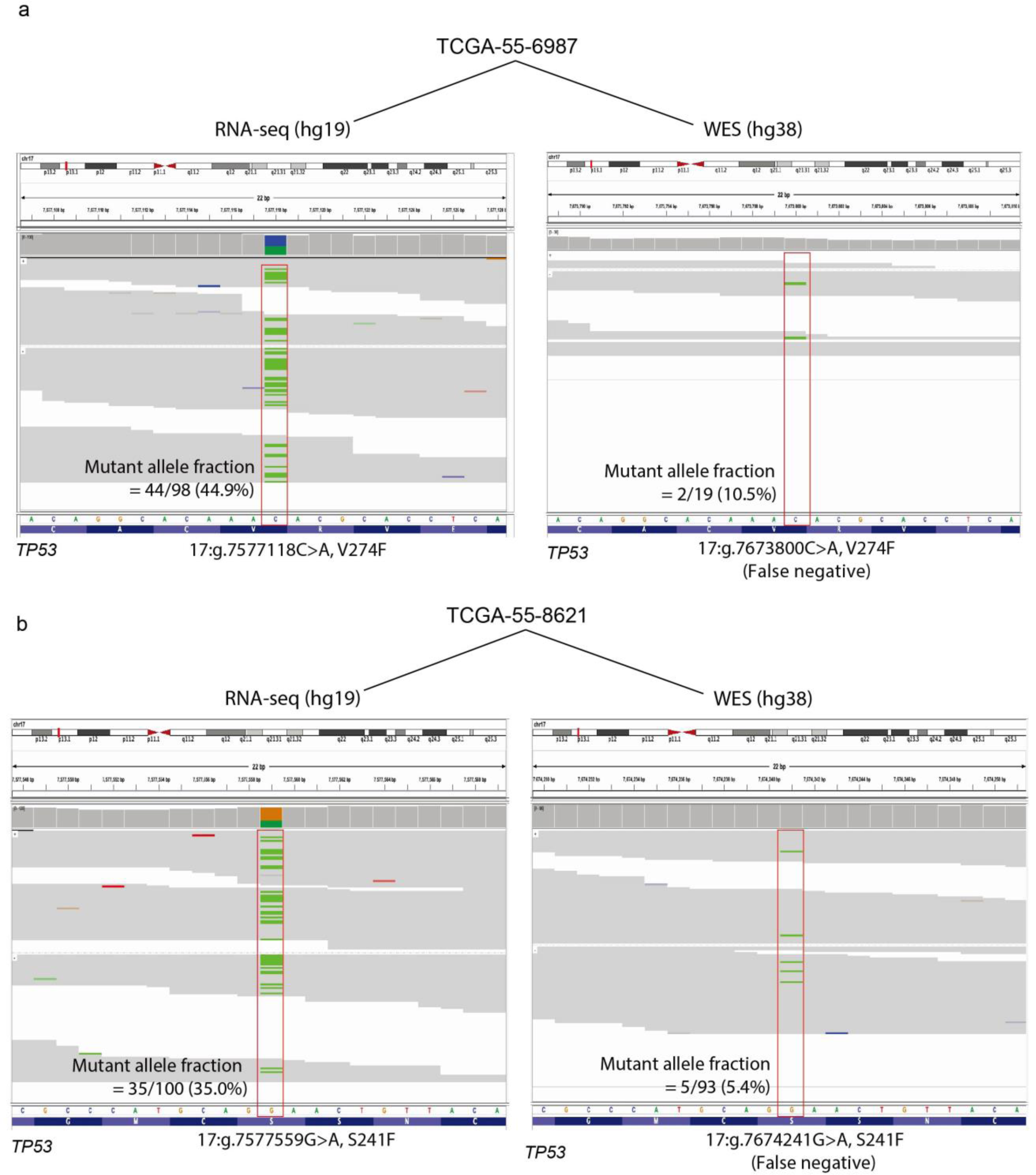
Examples of *TP53* somatic mutations identified from RNA-seq data but missed by TCGA WES calls. Mutant allele fraction (MAF) is measured by the ratio between the “number of reads supporting the mutant allele” and the “total number of reads” covering the mutation site.

**Supplementary Figure 9.**
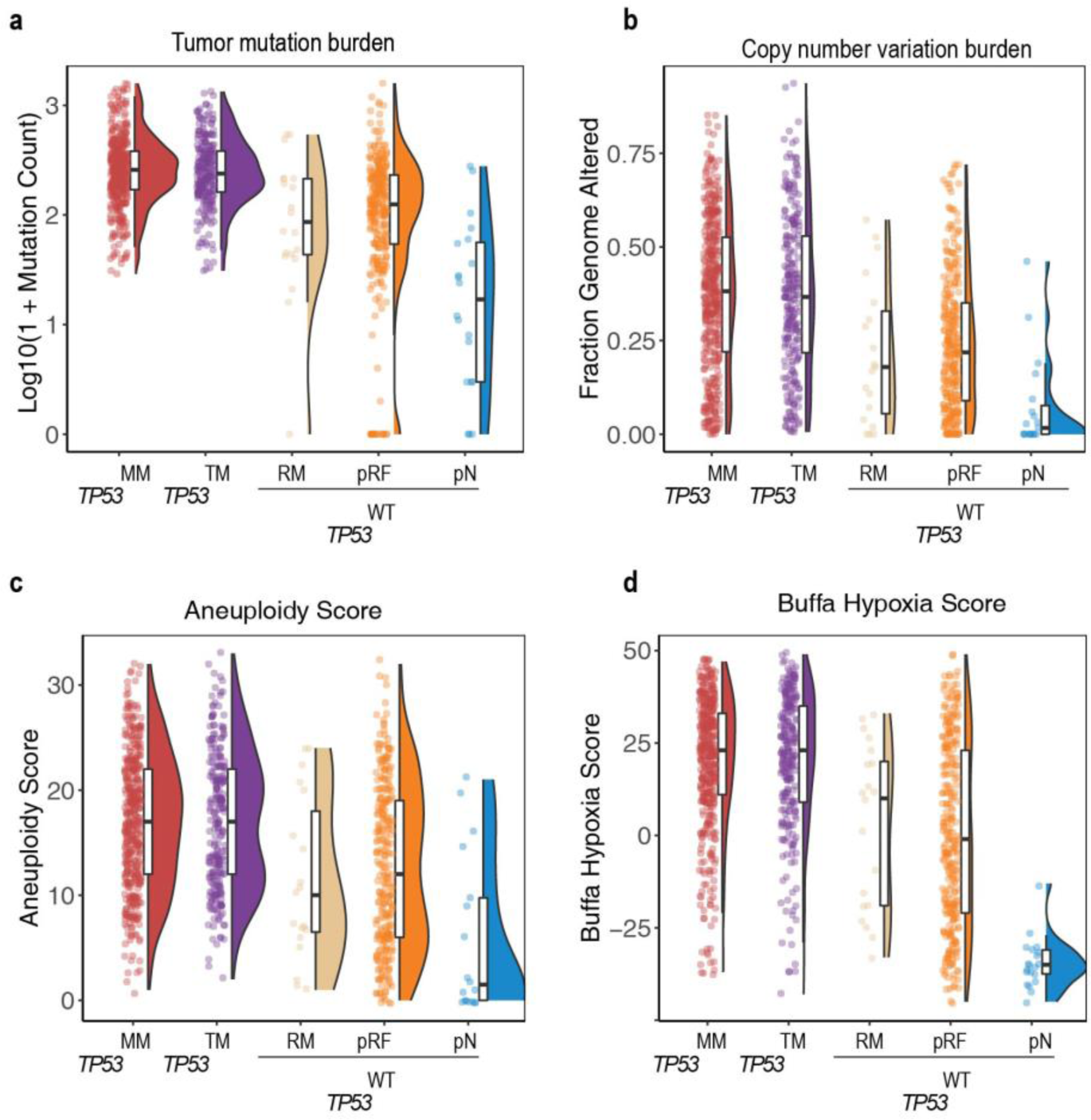
p53 mutants identified from RNA-seq data (indicated as the RM group) exhibit similar genomic characteristics as the *TP53*^MM^ and *TP53*^TM^ groups. Compared to the *TP53*^WT^-pN tumors, tumors of the RM group show increased tumor mutation burden (**a**), copy number variation burden (**b**), aneuploidy score (**c**), and Buffa hypoxia score (**d**).

## Supplementary tables

**Supplementary Table 1 List of p53 target genes used in this study**

**Supplementary Table 2 CESs of p53tru-DR and p53tru-UR genes in LUNG, BRCA, COAD, HNSC, STAD, UCEC, LIHC and Pan-cancer (including 9 types or subtypes) cohorts**

**Supplementary Table 3 The performance metrics of the SVM model in TCGA LUNG, BRCA, COAD, HNSC, STAD, UCEC, LIHC and Pan-cancer cohorts**

**Supplementary Table 4 List of SVM training samples, prediction samples, *TP53* original genetic states, SVM-predicted states and probabilities of TCGA cancers analyzed in our study**

**Supplementary Table 5 Genomic and clinical characteristics including “Mutation Count”, “Fraction Genome Altered”, “Aneuploidy Score”, “Buffa Hypoxia Score”, “overall survival (OS) time (day)”, and “OS status” of TCGA LUNG, BRCA, COAD, HNSC, LIHC, STAD, UCEC and Pan-cancer cohorts**

**Supplementary Table 6 Gene signatures reflecting chemo- or radiosensitivity**

**Supplementary Table 7 Median survival days and CES values of PDX models treated with placebo and radiation therapy (RT)**

**Supplementary Table 8 List of lung and breast cancer samples that have TP53 missense mutations detected from RNA-seq**

**Supplementary Table 9 Summary of SVM models according to the DOME (Data, Optimization, Model and Evaluation) recommendations**

## Notes

### Competing Interest Statement

The authors have declared no competing interest.

### Summary of Updates

The main changes are: 1.Included an independent LUAD dataset to indicate that p53-regulated genes can effectively reflect the functional status of p53 (page 5, line 128-137; page 15, line 465-472); 2. Performed Fisher's exact test and KEGG enrichment analysis to indicate that although there is potential involvement of other factors in influencing p53-regulated genes, TP53WT-pRF samples as well as the p53-regulated genes are closely related to TP53 (page 12, line 359-372); 3. Evaluated potential contributions of p53-regulated genes and p53 functional partners, to dissect predicted TP53WT-pRF (page 13, line 384-389); 4. Examined the treatment response of TP53WT-pRF samples using drug imputation data of TCGA patients, and discussed the results (page 13, line 390-400); 5. Modified the subheadings in the Result section to make them more informative, numbered the equations, made formatting changes according to the journal requirements, and made revisions throughout the manuscript to enhance clarity.

